# An *in vitro* neurogenetics platform for precision disease modeling in the mouse

**DOI:** 10.1101/2022.01.21.477242

**Authors:** Daniel E. Cortes, Mélanie Escudero, Austin C. Korgan, Alyssa Edwards, Selcan C. Aydin, Steven C. Munger, Kevin Charland, Zhong-Wei Zhang, Kristen M.S. O’Connell, Laura G. Reinholdt, Martin F. Pera

## Abstract

The power and scope of disease modeling can be markedly enhanced through the incorporation of broad genetic diversity. In the the mouse, the introduction of pathogenic mutations into a single inbred strain sometimes fails to mimic human disease. We describe a cross-species precision disease modeling platform that exploits mouse genetic diversity to bridge cell-based modeling with whole organism analysis. We developed a universal protocol that permitted robust and reproducible neural differentiation of genetically diverse human and mouse pluripotent stem cell lines, then carried out a proof of concept study of the neurodevelopmental gene *DYRK1A*. The results *in vitro* reliably predicted the effects of genetic background on *DYRK1A* loss of function phenotypes and identified optimal mouse strains for *in vivo* disease modeling. Transcriptomic comparison of responsive and unresponsive strains identified novel molecular pathways conferring sensitivity or resilience to *DYRK1A* loss, and highlighted differential mRNA isoform usage as an important genetic determinant of response This cross-species strategy will be an powerful tool in the functional analysis of candidate disease variants identified through human genetic studies.

Teaser: Genetically diverse mouse embryonic stem cells provide a rapid approach to precision modeling of human disease.

## Introduction

There is an increasing awareness of the importance of genetic diversity in cellular disease modeling, both for scientific reasons, and in relation to the issue of equitable access to biomedical innovations arising from *in vitro* discovery platforms^1^. The scientific benefits of genetic diversity in disease modeling extend to studies *in vivo*. Most mouse models for heritable human disorders have been created in a few commonly used inbred strains. However, genetic background can have profound effects on the phenotype of a particular mouse model. Thus, creating a gene knockout or mutation model on a single genetic background may or may not yield a phenotype relevant to human disease. This impediment to disease modeling has long been apparent from studies of mouse knockouts on different genetic backgrounds^2,3^, and can generally be related to the effects of modifier genes^4^.

To a considerable degree, incorporating greater genetic diversity into disease modeling in the mouse can provide a more reliable reflection of human disease pathogenesis than studies with a few inbred strains. A number of recent reports have shown clearly how the use of genetically diverse strain backgrounds can enhance our ability to model human sensitivity and resilience to disease, as exemplified in studies of Alzheimer’s Disease, Type 2 diabetes, autism and gastric cancer^5–12^. However, the establishment, maintenance and use of mouse diversity panels requires significant infrastructure, animal husbandry, and genetics resources, limiting their implementation for many investigators.

In recent years, mouse embryonic stem cell (mESC) lines from genetically diverse mouse strains have become available, and they have been used *in vitro* to study the genetic basis of the permissiveness of strains for mESC establishment, the regulation of the pluripotent state and early lineage specification, and the stability of genomic imprinting *in vitro*^10–17^. To apply these approaches in disease modeling, it is important that differentiation protocols provide for similar high yields of relatively pure cell populations across genetically diverse cell lines. Here we report a novel paradigm for the application of genetically diverse mESC lines to *in vitro* disease modeling in the central nervous system (CNS), a precision neurogenetics approach that has the potential to make mouse genetic diversity more accessible to the research community, and to provide for facile selection of appropriate mouse genetic backgrounds for human disease modeling.

Experimental platforms based on human pluripotent stem cells (hPSC), including two-dimensional cultures and three-dimensional organoid models of specific brain regions, are playing an increasing role in the study of the development of the central nervous system, and in unraveling the genetics of brain disorders^18^. However, most neurologic or psychiatric disorders are highly complex, involving the interaction of multiple systems inside and outside of the CNS and behavioral phenotypes that cannot be observed *in vitro*. We aimed to provide a facile link between human disease models and mouse diversity genetics, to enable precision disease modeling. We first established a set of robust universal neural differentiation protocols that enable production of key CNS cells of interest from genetically diverse mouse inbred strains and hPSC. These protocols are applicable across mESC and hPSC representative of wide genetic diversity, encompass multiple stages of neural development, and provide for functional readouts. Using high content screening, RNA-seq, electrophysiology, axonal regeneration paradigms, and knockout models, we provide a proof-of-concept study in the form of a chemogenomic and genetic analysis of the effects of loss of the neurodevelopmental gene, dual-specificity tyrosine phosphorylation-regulated kinase 1A (*Dyrk1a),* at multiple stages of neurogenesis and axonal repair. *Dyrk1a* was first discovered in Drosophila and the gene named “minibrain” because mutations in the gene causes size reduction of the optic lobes and central brain hemispheres^19^. *Dyrk1a* gene dosage affects many stages of neural development. In human, *DYRK1A* is located on Chromosome 21 and it has been associated with a range of conditions including Alzheimer’s Disease, Down syndrome, microcephaly, autism, and intellectual disability^20–25^. Patients harboring the same point mutation in the *DYRK1A* gene display different phenotypes ranging from severe autism to relatively minor neurological impairment^24^, strongly suggesting that the genetic background modifies the penetrance of the mutation.

Using genetically diverse mESC, we demonstrate the impact of genetic background on the outcome of DYRK1A inhibition or loss across multiple stages of neural development. With this approach, matching *in vitro* phenotypes in hPSC-based disease models with mESC models exhibiting similar sensitivity to pathogenic mutations enables rapid selection of mouse genetic backgrounds for *in vivo* modeling studies, and will provide a platform for prospective analysis of the power of *in vitro* mutant phenotyping to predict disease phenotype *in vivo,* a critical component of functional genomics analysis going forward.

## Results

To achieve our objectives in precision disease modeling, we had to develop and validate *in vitro* methodology for the assessment of the effects of genetic variants on neural development and disease across eight highly genetically diverse mESC representing the founders of the Collaborative Cross (CC founders, A/J (AJ), C57BL/6J (B6), 129S1/SvlmJ (129), NOD/LtJ (NOD), NZOHiLtJ (NZO), CAST/EiJ (CAST), PWD/PhJ (PWD, in place of the closely related PWK/PhJ), WSB/EiJ (WSB)), and to carry out cross-species comparisons with hPSC. First, we developed universal differentiation protocols that functioned efficiently, then we tested the validity of our in vitro platform for its ability to predict how individual genotypes would respond to disease mutations.

Initially we trialed widely-cited differentiation protocols used to generate neurons from mESC^26–31^ across a genetically diverse stem cell panel. These protocols failed when applied to strains other than 129. Therefore we developed the protocol shown in Fig S1a. With mESC from the B6 strain, by day 8 abundant neurites and neural rosettes are visible (Fig. S1b), and cells were passaged for expansion using EGF and FGF2 for three days. Decay in pluripotency markers and increase in neural markers were revealed by qPCR (Supplementary Figure 1c). At the end of this period, cells could be frozen, or passaged for further maturation by the withdrawal of mitogens and the addition of growth factors.

After the withdrawal of mitogens, NPCs exited the cell cycle and started to generate more elongated processes (Figure 1a). The timing of differentiation was similar across all strains (Fig. S2). After 7 days of differentiation, neurons represented around 90% of the overall cell population among all strains studied (Figure 1b,c). Among these neurons, glutamatergic cells represented around 75% of the neuronal population (Figure 1,d,e). Practically all neurons expressed the forebrain marker FoxG1 and cells expressing all individual six cortical layer markers were observed, across all strains analyzed strains at similar frequencies (Figure1b,e). We also analyzed the electrophysiological activity using whole cell patch clamp. Evoked action potentials were elicited as well as spontaneous activity, both appearing in cultures of all strains after 2 weeks of differentiation (Figure 1f) and coinciding with the presence of pre and post synaptic proteins (Figure 1d). Using multielectrode arrays (MEA) we found spontaneous field potentials (Figure 1g, upper panel) and network activity in all strains (Figure 1g, lower panel). We could also produce cortical organoids that displayed spontaneous calcium signaling (Figure 1h, i).

**Figure 1.**
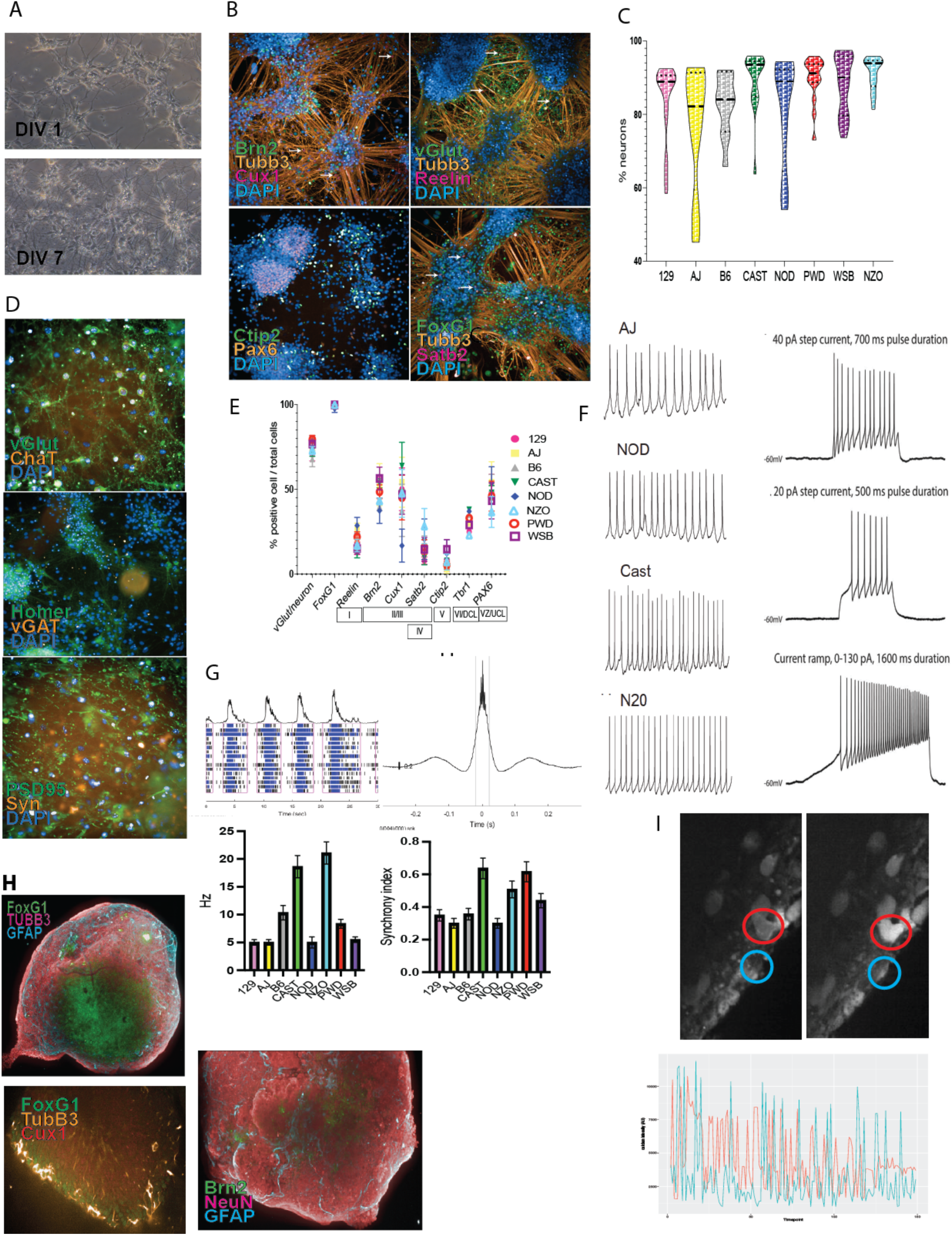
mESC-derived NPC follow the default differentiation pathway into cortical neurons and cortical cerebral organoids. a) Neurons start to develop processes one day after seeding and they become more convoluted after 7 days. b) Immunostaining for different cortical layer markers. c) Violin plot showing the percentage of mESC-derived neurons for each of the eight different mESC strains studied. d) Immunostaining for neurotransmitters and synaptic markers. e) Plot showing glutamatergic, forebrain and cortical layer markers from all the strains of mESC-derived neurons. Data are represented as mean ± SEM. f) Representative images of whole cell patch clamp neuron recordings, spontaneous activity on the left and evoked potentials on the right. g) representative recordings of mESC-derived neurons on MEA plates. Top left is a raster plot. Top right is a representative image of synchrony within the well. Bottom left, column plot showing spike activity (Hz) among the neurons derived from all strains. Bottom right shows the synchrony index among the neurons derived from all strains. Data are represented as mean ± SEM. h) Representative images of mESC-derived organoids showing different cortical markers. i) Top panel, calcium imaging of mESC cerebral organoids. Circles show two examples of cells with changes in calcium activity. Bottom panel, plot showing the activity of the two cells in the upper panel followed through time.

Modifications to the protocol, using morphogens to achieve patterning, enabled us to generate motor neurons, GABAergic neurons, and dopaminergic neurons from all strains tested at similar efficiencies (Figure 2). We reasoned that by adjusting the developmental timing between mouse and human cultures (Figure 3a), our protocol could be adapted to differentiate neurons from hPSC of diverse genetic backgrounds. We tested 10 genetically diverse hPSC from the iPSC Consortium for Omics Research (iPSCORE) representing equally both sexes and diverse ancestries. Once again, we were able to obtain neural progenitors and cortical neurons, and we found that all cell lines differentiated with similar efficiencies regardless of their genetic background (Figure 3b,c); we observed spontaneous synchronous electrical activity in these cultures (Figure 3d). With appropriate patterning (Figure 3e) we obtained motor neurons that also showed network activity (Figure 3f,g), as well as cortical and midbrain dopaminergic organoids that displayed spontaneous calcium signaling (Fig. S3,4 and Movie S1).

**Figure 2.**
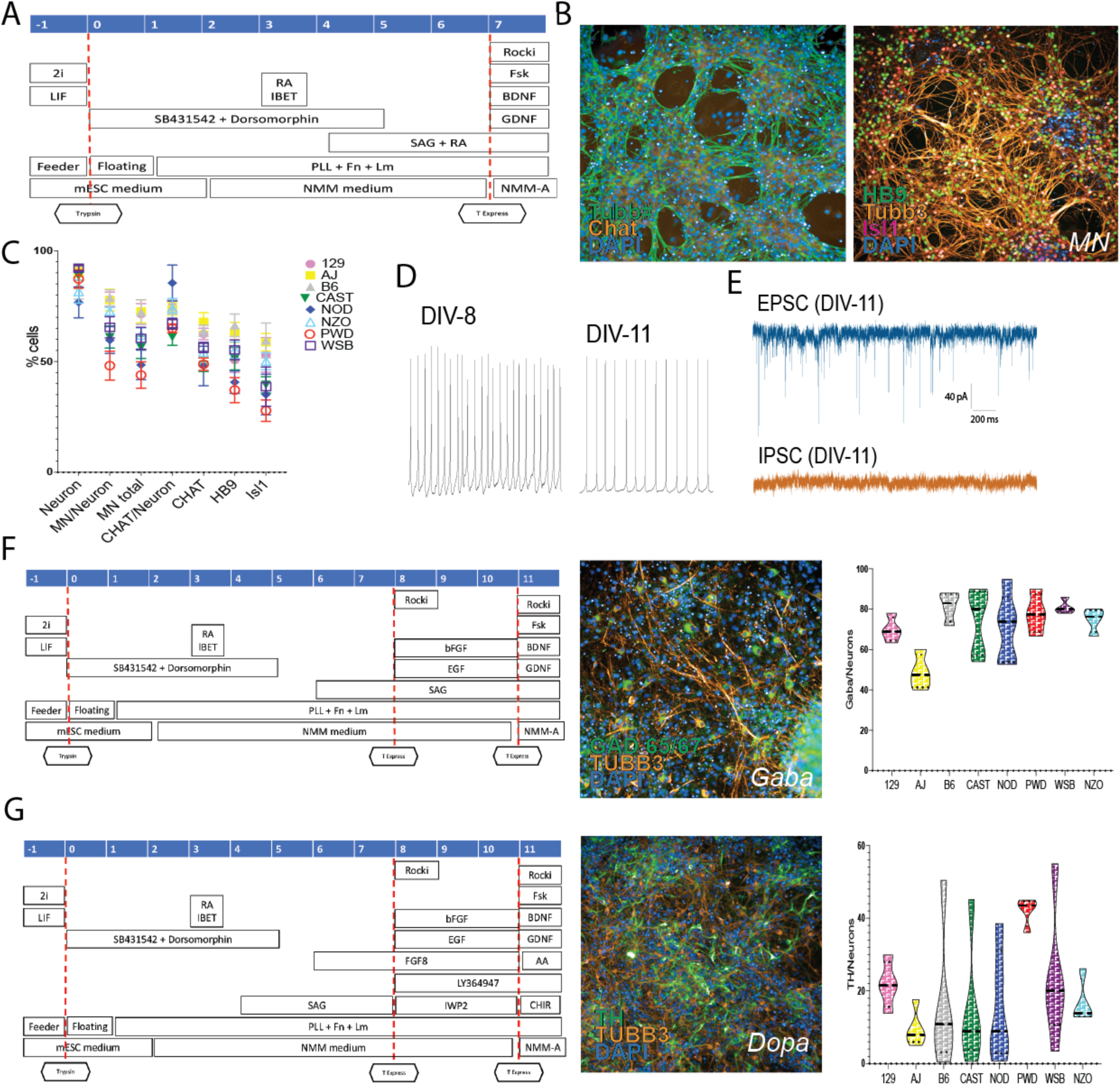
Generation of specific neuronal subtypes by molecular patterning. a) Scheme of the differentiation protocol to generate mESC-derived MN. b) Immunostaining for MN markers. c) Plot showing the differentiation efficiency among the eight mESC strains. Data are represented as mean ± SEM. d) Whole cell patch clamp recording performed at day 8 (left) and 11 (right) of the MN differentiation. e) Representative EPSC and IPSC recordings on mESC-derived MN. f) Scheme of the differentiation protocol to generate mESC-derived GABAergic neurons (left) with a representative image of GABAergic neurons (center) and quantification of the differentiation efficiency among all mESC tested (right). g) Scheme of the differentiation protocol to generate mESC-derived dopaminergic neurons (left) with an image of dopaminergic neurons (center) and quantification of the differentiation efficiency among all mESCs tested (right).

**Figure 3.**
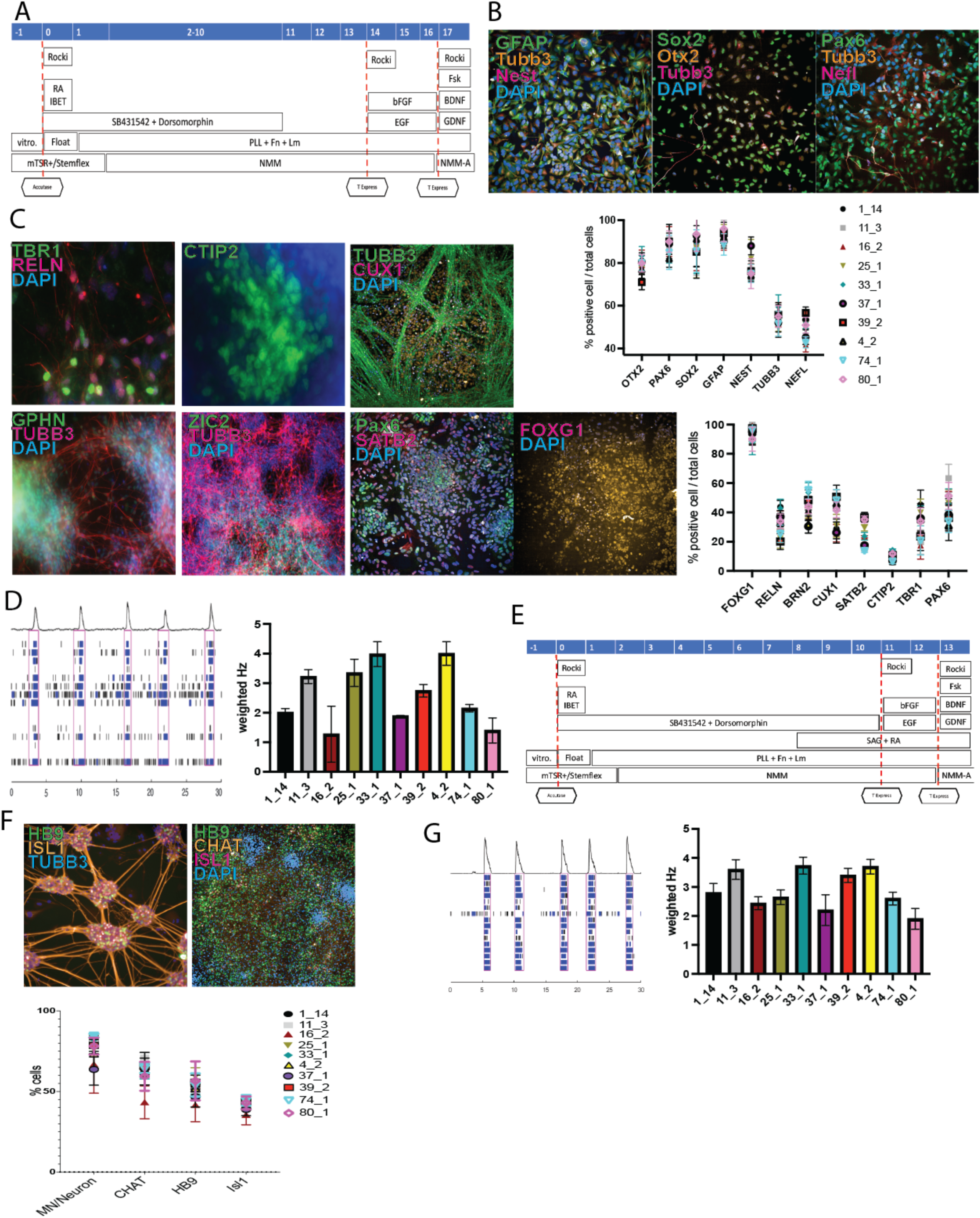
Genetically diverse hPSC can be differentiated into cortical and MN. a) Scheme of the differentiation protocol to generate hPSC-derived cortical neurons. b) Immunostaining of hPSC-derived NPC and quantification of NPC markers (bottom). Data are represented as mean ± SEM. c) Cortical layer markers after 3 weeks of maturation and quantification (right). Data are represented as mean ± SEM. d) Representative raster plot of hPSC-derived cortical neurons obtained using an MEA plate after 3 weeks of maturation (left). Quantification of frequency among the 10 genetically diverse iPSCs (right). e) Scheme of the differentiation protocol to generate HPSC-derived MN. f) Representative images of hPSC-differentiated MN showing phenotype-related markers (top) and quantification efficiency among the 10 genetically diverse iPSCs (bottom). g) Representative raster plot of hPSC-derived MNs obtained using an MEA plate after 3 weeks of maturation (left) and quantification of frequency among the 10 genetically diverse iPSCs (right).

Having developed this differentiation platform, we proceeded to conduct a proof-of-concept study to examining how genetic background impacts on the effect of DYRK1A inhibition during neuronal development at different stages. *Dyrk1a* also regulates axonal growth and guidance, so we decided to assess an experimental model of axon regrowth as well. In most studies, we used the well-characterized DYRK1A inhibitor ID-8; this inhibitor is highly selective against DYRK1A compared to other kinases. ID-8 IC_50_ values for DYRK1A, GSK3b, and CLK1 are 78, 450, and 4200 nM respectively, and DYRK2, DYRK3, and DYRK4 are not affected at all^32–35^.

To ascertain that differences in phenotypes observed in response to DYRK1a inhibition or haploinsufficiency were not merely a consequence of differences in gene coding sequences (and by extension protein structure) or expression levels across the strains, we examined both (Fig. S5). Despite some divergence in the gene sequences, notably for CAST and PWD (Fig. S5a), the SNV do not affect the protein sequence and all variants are synonymous (Supplementary Table 1). Examination of expression quantitative trait loci (eQTL), in mESC^14,36^ revealed a local eQTL in CAST and PWD/K (Fig. S5b). Nevertheless, we could not find any protein QTL among all the strains in mESC, and direct measurement of protein levels in mESC revealed that although there is a slight decrease of DYRK1A protein in CAST, NOD, and PWD, these differences were not statistically significant (Fig. S5c,d,e and for neural progenitors, Figure 6 below).

### Effects of DYRK1A inhibition on mESC maintenance and lineage specification vary across genetically diverse mouse strains

It was previously shown that DYRK1A inhibition by ID-8 in mESC and hPSC helps to maintain the pluripotent state, and that either chemical inhibition or knockdown hampers specification from the pluripotent state to the neural lineage in hPSC^32,35^. In order to study pluripotency maintenance and lineage biases across mESC from different strains to compare with previous work on 129 mESC and hPSC, we cultured the eight mESC lines in media supplemented with four combinations of factors: LIF+2i (LIF plus PD0325901 and CHIR99021,optimal maintenance of naïve state mESC,^37^), LIF+ID-8, LIF, and ID-8. These cell lines were previously shown to vary in their response to LIF and in the stability of the undifferentiated state^14,38^; some lines remain in the undifferentiated state and others switch on lineage markers as soon as LIF+2i optimal conditions are abandoned, especially those mESC lines from strains regarded to be recalcitrant to stem cell derivation^13^.

Using high content screening (HCS), we analyzed the expression of ten different pluripotency and lineage specific markers during four days of culture in the above conditions (Figure 4a,b). Only LIF+2i was able to maintain the expression of the pluripotency associated factors POU5F1, NANOG, and SOX2 in all cell lines; nonetheless, PWD and NZO expressed these markers at low levels under all conditions.

**Figure 4.**
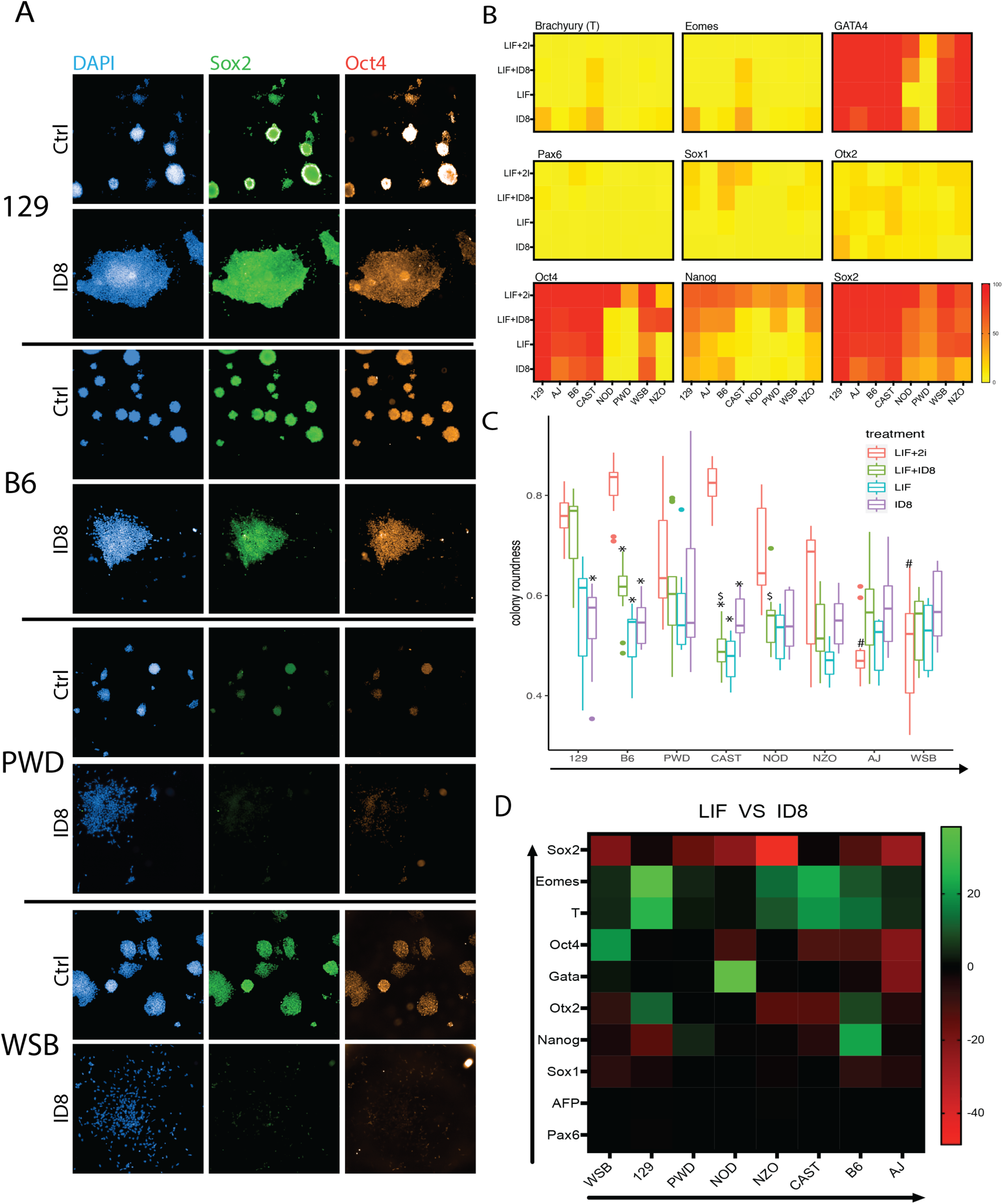
Effects of DYRK1A inhibition on mESC maintenance and lineage specification vary across genetically diverse mouse strains. a) Immunofluorescence images showing immunostaining for pluripotency markers in the eight different CC founder mESC cultured in four different culture conditions for four days. b) Heatmaps showing the percentage of cells expressing the markers in the mESC as measured by HCS. c) Morphometry analysis of the mESC colonies. d) Heatmap comparing the effects of LIF vs ID-8 treatment on the overall proportion of cells expressing the analyzed markers in the eight different mESCs, determined by HCS. Red indicates decreased percentage of cells expressing the associated marker in ID-8 compared to LIF, green means higher expression. Cell lines and markers are arranged according to the overall degree of change (arrows). *, represents statistically significant differences against the control condition (LIF+2i) within the same cell line. #, represents statistically significant differences among cell lines treated with 2i+LIF and compared to the standard cell line 129. $, represents statistically significant differences among cell lines treated with 2i+ID8 and compared to the standard cell line 129. &, represents statistically significant differences among cell lines treated with LIF and compared to the standard cell line 129. %, represents statistically significant differences among cell lines treated with ID-8 and compared to the standard cell line 129 where a significant difference is indicated when p< 0.05. Means +/-SEM are shown. Kruskal-Wallis and Dunn’s test were conducted. n = 4 biological replicates.

ID-8 was originally found to substitute for LIF in mESC maintenance on the 129 background ^32^. Comparing cultures grown in LIF+2i with ID-8, we note that 129, AJ, and B6 show the best maintenance of NANOG expression in ID8 alone (Figure 4b, bottom panel). We also examined the proportion of cells expressing markers of the three embryonic germ layers (Figure 4b, top two panels). Those cell lines that showed consistent expression of pluripotency markers in ID-8 also showed low expression of most germ layer lineage markers. Colony morphometry using HCS provides another indicator of cellular response to the culture environment. Loss of stem cell marker expression generally correlated with changes in colony morphology, as morphometry showed that cell colonies became less compact and circular in ID-8 alone, and LIF+ID-8 enhanced roundness in 129 only (Figure 4c). Comparing LIF vs. ID-8 alone, the two conditions maintain pluripotency associated markers to a similar degree, but in 129, B6 and CAST, mesodermal markers are expressed at higher levels (Figure 4b,d). In summary these results confirm previous studies in 129 mESC and hPSC indicating that DYRK1A inhibition can help maintain the undifferentiated state. AJ and B6 derived mESC respond to ID-8 similarly to 129, but strains that are poorly maintained in LIF+2i are not rescued by DYRK1A inhibition.

### DYRK1A inhibition differentially blocks neural progenitor proliferation in a strain-dependent fashion

Next, we used our differentiation system to study multiple stages in neural development, from neural lineage specification through to neurogenesis, neuronal differentiation, and maturation. For this, we differentiated the eight mESC lines into NPCs using our newly developed protocol in the absence or presence of ID-8, and analyzed the cultures at the beginning and on days two, five, and eight via immunostaining and HCS (Figure 5a,b). Following induction of neural specification in the presence of ID-8, B6 cultures contained lower percentages of OTX2 and TUBB3 positive cells, and displayed a marked reduction in the proportion of KI67 positive cells compared to control. Inhibition of the proliferation of neural progenitor cells during development is a hallmark of *Dyrk1a* haploinsufficiency. Further examination of the proliferation of neural progenitors from additional mESC clones of B6, AJ, 129 and WSB using KI67 and MCM2 immunostaining confirmed that the proliferation of B6 cells was strongly inhibited by ID8, whereas the latter three strains were unaffected (Figure 5c,d).

**Figure 5.**
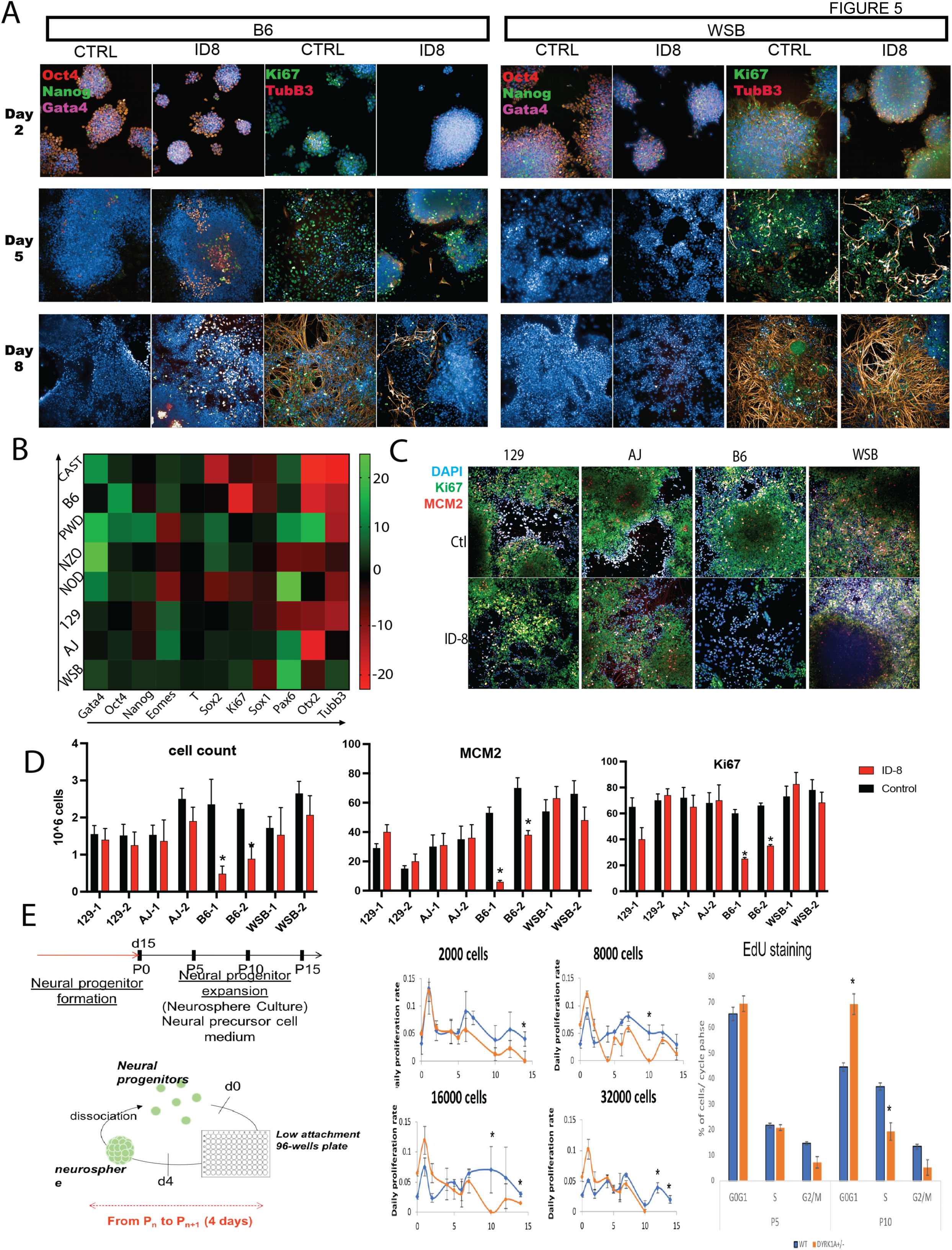
DYRK1A inhibition differentially blocks neural progenitor proliferation in a strain-dependent fashion. a) Representative images of the B6 and WSB mESC lines differentiated into NPC in the presence of the DYRK1A inhibitor ID-8 and analyzed at day 2, 5 and 8. b) Comparison between the proportion of cells expressing the indicated markers as measured by HCS in control and ID-8 treated cells after eight days of differentiation. Red indicates decreased percentage of cells expressing the associated marker in ID-8-treated cells compared to control whereas green means an increase. Cell lines and markers are arranged according to the overall degree of change (arrows) and n = 4 biological replicates. c and d) Images and plots representing the cell count of proliferating NPCs derived from six independent cell lines different from those used in b. NPCs were cultured with or without ID-8 for 5 passages. B6 showed a decreased number of cells and proliferative markers (Ki67 and MCM2) compared to its control whereas 129, AJ and, WSB were not affected. n = 4. * Statistically different from its control, p < 0.05. Kruskal-Wallis and Dunn’s test were conducted e) Long-term proliferation assay in hPSC-derived NPC. Left, experimental design (cells were kept in proliferative conditions as neurospheres for 15 passages (2 months total); middle, cell numbers over time; right. percentage of cells in each cycle phase measured by EdU staining in WT and DYRK1A+/-. Experiment shows DYRK1A +/-NPC undergo quiescence faster than control. Two-tailed T test, n = 4, * = p < 0.05.

We previously observed an inhibition of neural specification in hPSC similar to that noted here^35^. We therefore examined the effect of heterozygous loss of *DYRK1A* on neural progenitor proliferation in the human induced pluripotent stem cell (hiPSC) line KOLF2.1J^39^. The susceptibility of B6 neural progenitor cells to inhibition of proliferation by ID-8 was mirrored in the reduced growth of *DYRK1A*+/-human iPSC neural progenitors relative to controls (Figure 5e). Thus, in terms of effects of DYRK1A inhibition on the maintenance of the undifferentiated state, interference with neural specification, and reduction of neural progenitor proliferation, the B6 mESC display the best face validity for modeling the response of human pluripotent stem cells to DYRK1A inhibition or haploinsufficiency.

To corroborate that the observed neural differentiation impairment is caused by ID-8-induced DYRK1A inhibition; we generated *Dyrk1a* knock-out mESC lines, specifically on B6, 129 and WSB backgrounds. We conducted NPC differentiation on these cell lines and analyzed the phenotype. Similar to results with chemical knockdown of DYRK1A, we observed that the only strain severely affected by *Dyrk1a* deletion was B6, whilst neural progenitor specification, proliferation and differentiation in WSB and 129 cell lines were indistinguishable from parental controls (Figure 6a,b). During B6 knockout cell line differentiation, we observed a decreased expression of proliferation and neural markers, and a decreased number of cells (Figure 6a,b). The proportion of cells expressing DYRK1A, and the intensity of staining, was similar in wild-type cells of all three strains (Figure 6b).

**Figure 6.**
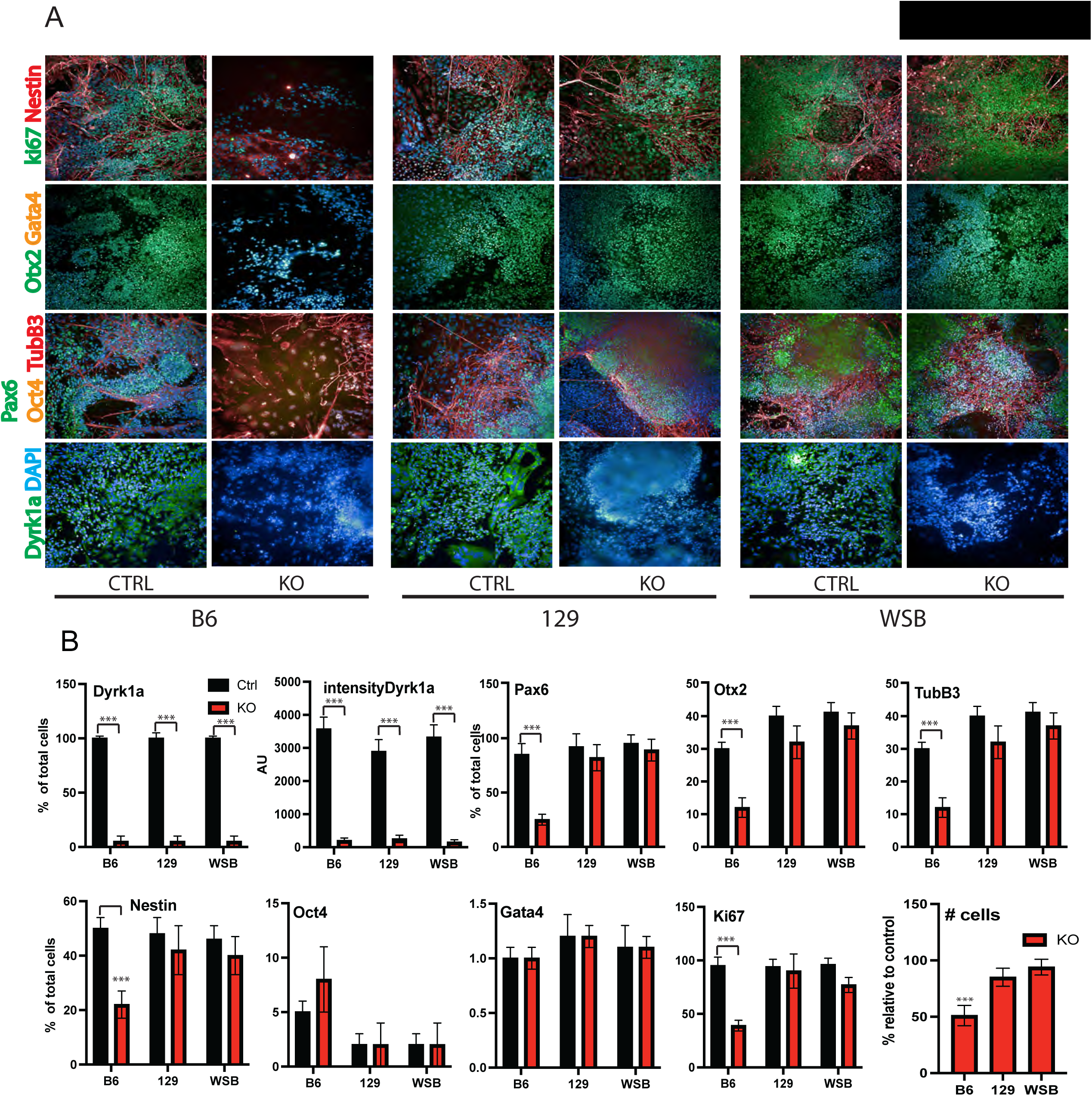
Neural differentiation of WT and Dyrk1a-knockout mESC. a) Representative images at day eight of NPC differentiation showing robust differentiation in 129 and WSB but B6 *Dyrk1a*-knockout. b) Quantitation of proportion of cells expressing DYRK1A (and average intensity of DYRK1A staining) and markers of neural differentiation, pluripotency, and proliferation, measured by immunostaining and high content screening in the three strains. Kruskal-Wallis and Dunn’s test were conducted. * = p < 0.05 compared to its own control n = 3 biological replicates.

Although all strains had different degrees of response to ID-8 during neural progenitor specification and expansion, WSB was the most unresponsive while B6 was the most affected. Bulk RNA-seq was performed on these two cell lines at day eight of differentiation to identify the differences in gene expression associated with the extreme phenotypic differences found between WSB and B6. This difference in response to the inhibitor is evident from the numbers of differentially expressed genes (DEG); there were more than 700 DEG between B6 control and ID-8 treated cultures, but only 13 for control versus treated WSB (Figure 7a).

**Figure 7.**
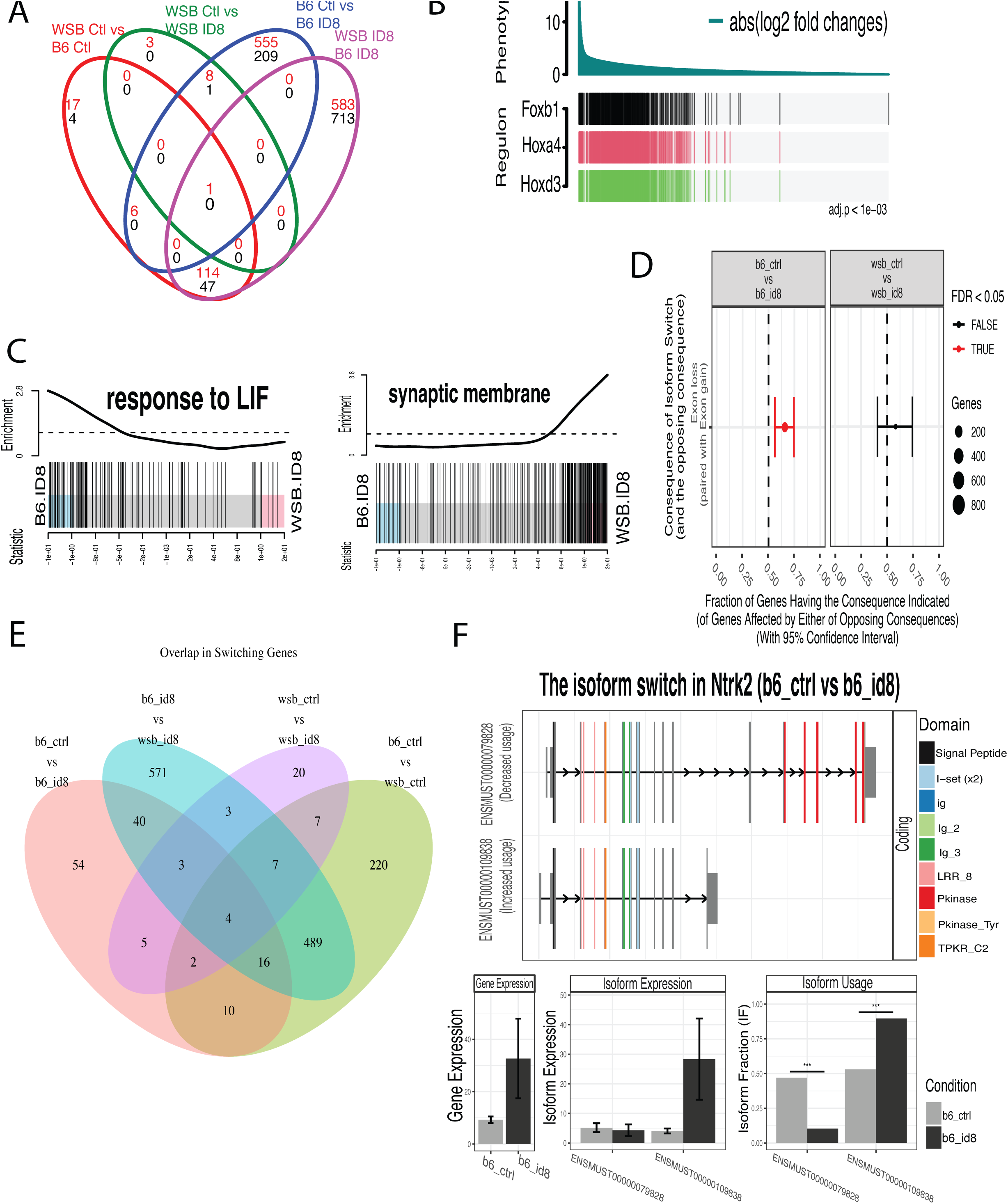
RNA-seq analysis reveals the genes associated with the impairment in neural induction in B6-derived NPCs. a) Venn diagram of DEG for each set of comparisons. Number in red represents downregulated genes, black indicates upregulated gene. b) RTNA for the top regulons associated with differences in ID-8 treated WSB- and B6-derived NPCs. c) Barcode plots showing two exemplary GOs differentially affected in WSB and B6 when treated with ID-8. Genes are ranked from left to right by increasing log-fold change represented by the vertical bars. The curve shows the relative local enrichment of the bars in each part of the plot. d) Consequences of isoform switch showing a lost of exons in ID-8 treated B6 neurons. e) Venn diagram showing the global isoform switch differences in B6 vs WSB neurons when treated with ID-8. f) Switch in the *Ntrk2* isoform usage when B6 is treated with ID8, upper plot shows the color-coded domains present on each exon (bars). Bar plots indicating total gene expression, isoform expression and relative isoform usage proportional to the total gene expression.

Regulatory transcriptional network analysis (RTNA) revealed *Foxb1*, *Hoxa4*, and *Hoxd3* as the top regulons associated with the differences found between WSB and B6 in response to ID-8 treatment (Figure 7b). *Foxb1* is a regulator of neural progenitor differentiation ^40^, and is rapidly and transiently induced during cortical specification of hPSC ^41^. The two Hox genes are associated with hindbrain patterning in the CNS ^42^. These pathways, not previously associated with *Dyrk1a* loss, are consistent with interference in neurogenesis and specific perturbations in hindbrain development (reported below) associated with *Dyrk1a* haploinsufficiency.

Gene ontology (GO) analysis indicated that either under control conditions or in the presence of ID-8, DEG associated with the response to LIF were highly overrepresented in B6 vs. WSB (Figure 7c, left panel). The LIF signaling pathway is responsible not only for maintenance of mESC but also for maintenance of mouse neural stem cells *in vitro* and i*n vivo* ^43^. In the presence of ID-8, DEG associated with the GO term synaptic membrane were overrepresented in WSB relative to B6 (Figure 7c, right panel), consistent with progression of neural differentiation in the former but not the latter.

Using the RNA-seq data, we also analyzed gene isoform changes due to treatment and between B6 and WSB strains. Isoform usage can change considerably even when there is no overall change in gene expression, with functional consequences^44^, and *Dyrk1a* is known to regulate the activity of a number of RNA splicing proteins^45–48^. Global analysis showed that adding ID-8 induced a loss of exons in B6 strain but not in WSB (Figure 7d). We found 200 unique genes with alternative isoform usage when control B6 cells were compared to WSB cells; an increase of variant isoforms between the two strains was observed in the presence of ID-8. (Figure 7e). Of particular interest was the finding that *Ntrk2* underwent a change in isoform expression in B6 NPC when treated with ID-8, whereas in WSB these isoforms remained unchanged (Figure 7f). *Ntrk2* codes for a neurotrophic receptor that binds BDNF. We found that ID-8 treatment induces a switch from the canonical isoform to an isoform lacking the kinase domain. This isoform is unable to transduce signaling via tyrosine phosphorylation, and it is notable that BDNF signaling was implicated in the response to *Dyrk1a* haploinsufficiency in a recent proteomics study in B6 mice^49^. That study did not investigate alternative splicing but the results are compatible with our findings.

These results suggest that, in the B6 strain in which neurogenesis is affected, DYRK1A inhibition interferes with multiple pathways involved in neural specification, patterning, and differentiation. RNA-seq confirmed the resilience of WSB against, and susceptibility of B6 to, DYRK1A inhibition at the neural induction stage. Subsequent analysis highlighted key pathways in neurogenesis downstream of DYRK1A inhibition, and identified a profound effect of DYRK1A inhibition on mRNA isoform usage. In the case of BDNF signaling, our results implicated alteration of isoform usage as an important genetic determinant of the response of neural progenitors to chemical knockdown.

### DYRK1A inhibition delays functional maturation of neurons and modulates gene expression of maturation-related genes in a strain-dependent fashion

To understand how DYRK1A inhibition modulates physiological maturation of neurons, we used MEA plates capable of recording 16 field potentials per well on a 48-well plate to assess electrophysiological activity. We seeded young neurons and let them mature for six weeks while analyzing electrical activity at different time points (Figure 8a,b). Addition of ID-8 began two days after the NPCs were seeded and continued throughout. All cell lines were capable of generating action potentials and bursts of spikes, and creating networks, but not all cell lines matured at the same rate. CAST, B6, and 129 were the cell lines that matured the fastest, measured by Hz, bursting activity, and synchrony, while PWD took the longest (Figure 8a,b). We found that in most of the strains, addition of ID-8 inhibited the development of electrical activity; neurons fired less and bursts of activity were less frequent. Complementarily, interburst intervals, the relative silent time between two trains of activity, were longer for most cell lines when ID-8 was added. Quantitative analysis of a range of parameters at the peak point of control electrical activity (Figure 8b) confirmed inhibition of maturation across most strains by measurement of mean firing rate, number of bursts, prolongation of interburst intervals, and a reduction in spikes per burst. Two overall patterns were observed. In B6, CAST, A/J, and NOD, maturation was delayed and suppressed throughout the time course, whilst in PWD, 129, NZO, and WSB, maturation was delayed but activity recovered at the later time point (Figure 8b,c). Taken together, once again the functional maturation of neurons from B6 mESC was sensitive to ID-8, that of WSB mESC-derived cells were partially resistant to ID-8, and NZO derived neurons were the most resilient among all the strains. A decreased rate of functional network maturation is consistent with known effects of ID-8 on axonal growth and dendrite arborization.

**Figure 8.**
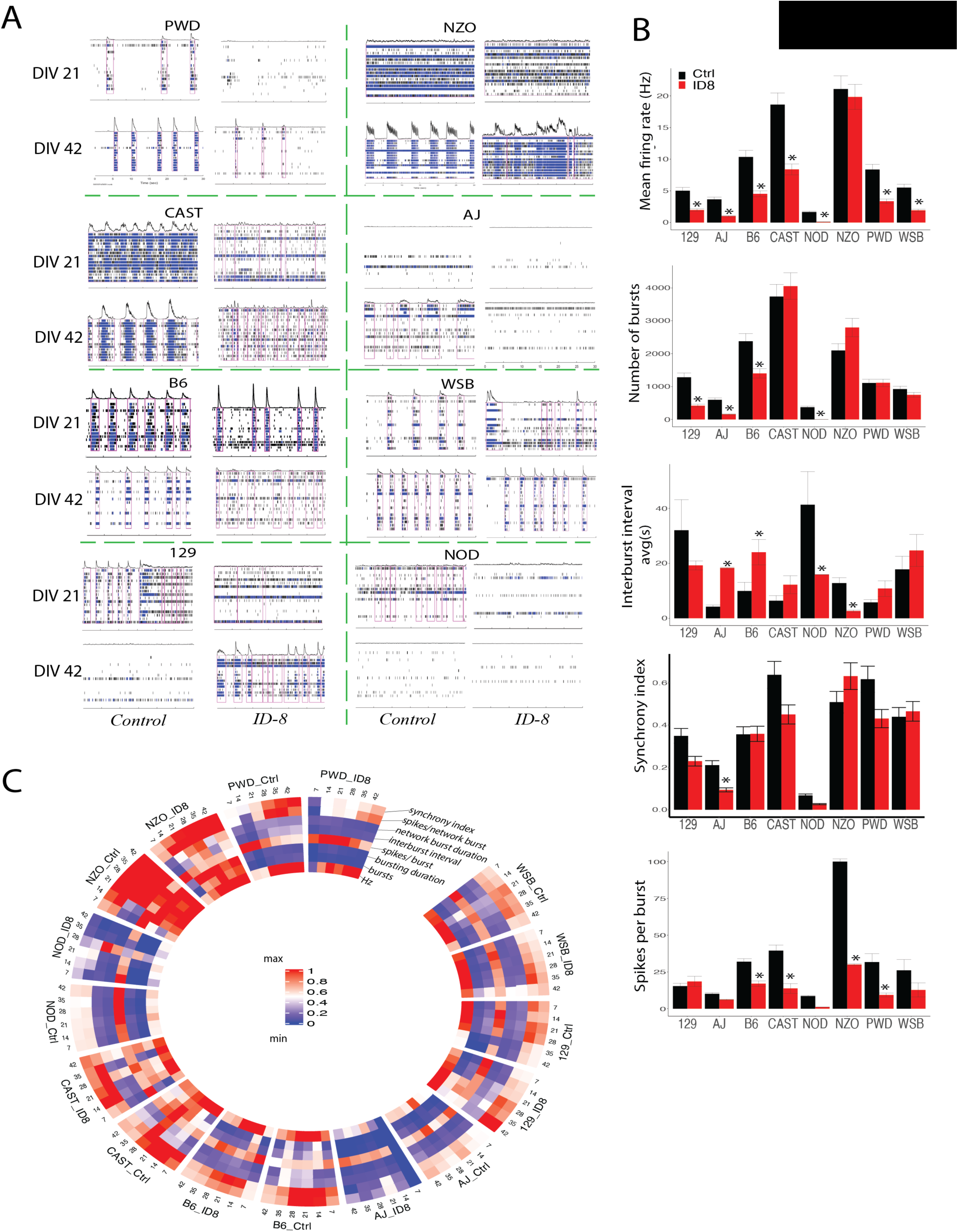
Effect of DYRK1A inhibition on delays in functional maturation of neurons by mESC strain. a) Examples of 30 second recordings of all differentiated neurons (21 and 42 days *in vitro*) in the presence or absence of ID-8. b) Plots for firing rate, number of bursts, interburst interval, synchrony, and spikes per burst. Because the neurons from different strains do not mature and survive equally, data form comparisons among the cell lines were taken at the timepoint where the highest frequency of firing (Hz) was recorded for each cell strain. Means +/-SEM are shown. Kruskal-Wallis and Dunn’s test were conducted. * = p < 0.05 compared to its own control n = 3 biological replicates. c) Circular plot showing different electrophysiological properties during 42 DIV in the presence of ID-8.

We next performed RNA-seq on WSB and B6 neurons. Analysis at this stage showed a similar pattern to the NPC stage on the MDS plot which separates the neurons by strain and treatment; it also showed more distance between B6 subgroups than the WSB counterpart indicating B6 was more affected by DYRK1A inhibition (Fig. S6a). After ID-8 treatment, there were 220 DEG between WSB and B6 (Fig. S6b). GO analysis of DEG differentially affected by ID-8 in the two strains identified biological adhesion and transmembrane receptor signaling, with WSB showing a greater enrichment following ID-8 treatment (Fig. S6c). Cellular adhesion molecules are important for neural development and axonal growth ^50^. *Rxrg*, *Dmrtb1*, and *lrf7* were the top regulons associated with the differences in response to DYRK1A inhibition between WSB and B6 neurons (Fig. S6d). *Rxrg* has been associated with neurodegeneration ^51^ and *Irf7* is a key mediator of the innate and adaptive immune responses ^52^ and positively regulates microglia polarization to the M2 state.

As we observed during neurogenesis, we also found more alternative splicing events in B6 compared to WSB neurons when treated with ID-8 (Fig. S6e). An example of this is *Slit2*, which exhibited isoform switches in B6 neurons when treated with ID-8 (Fig. S6f). *Slit2* is associated with axonal guidance and cell migration, and arborization^53^. All these developmental processes are affected by Dyrk1a mutations^54,55^.

### mESC strain differences in recovery from axonal injury in vitro reflect genetics of recovery from brain injury in vivo

There are marked strain differences in the ability of different mouse strains to respond to injury to the CNS ^56^. Abundant evidence confirms that *Dyrk1a* plays a role during neurite sprouting in normal development, and that gene dosage alteration affects axonal growth and dendrite arborization^23^, but it is not known whether these developmental roles of *Dyrk1a* might be recapitulated during axonal regeneration. As a further test of the predictive power of our *in vitro* system, we performed axotomy to analyze differences in axonal regrowth in neurons derived from mESC from CC founder strains.

We differentiated the mESC into neurons and cultured them into microfluidic chambers. (Figure 9a). On these devices, neurons were seeded on one chamber; axons could traverse microgrooves reaching into a second chamber whereas somas could not. After four days, axons grew through to the second chamber (Figure 9b). However, the number of axons growing through to the second chamber, and their length and density, indicated by the total area occupied by axons, varied amongst mESC strains (Figure 9c). At this point, axons were cut off and left to recover for four more days while treated or untreated with ID-8. In all cases, under control conditions, more axons grew across the device and had longer processes than before injury, except for B6 neurons (Figure 9b-d). CAST, NOD, and NZO grew significantly more following injury compared to all other strains. When ID-8 was added, most of the lines showed less growth than controls. WSB, AJ, and PWD axons occupied the same area after regrowth in the presence of ID8 as the control did (Figure 9d). WSB axons experienced the longest growth pre- and post-injury (both control and ID-8 treated). Notably, the extent of axon growth prior to injury was not predictive of the extent of regrowth post injury (e.g. 129 versus NZO versus B6, Figure 9c).

**Figure 9.**
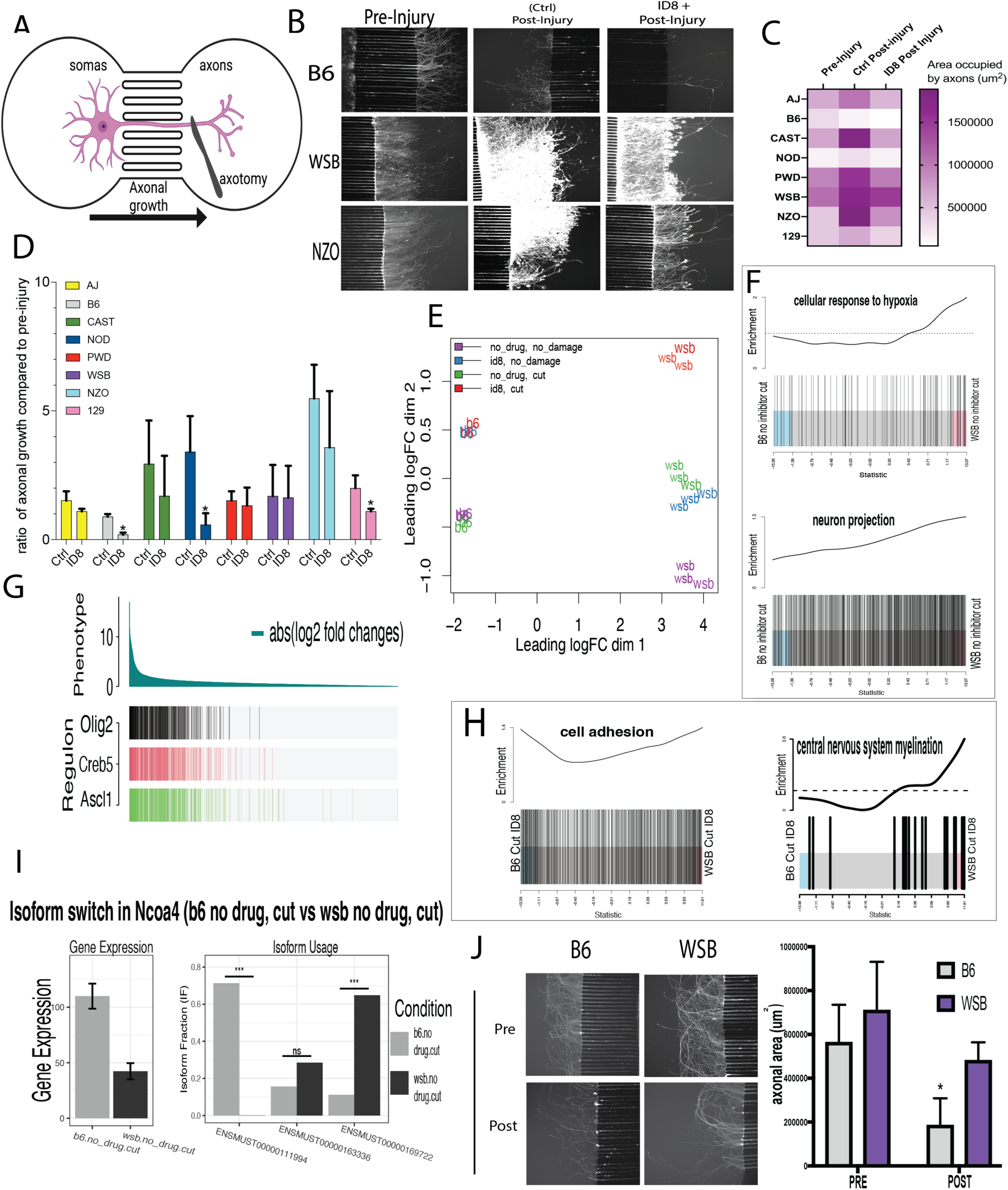
mESC derived neuronal strain differences in recovery from axon injury in vitro. a) Illustration of the microfluidic chamber used to isolate axons from somas. Image taken from chamber manufacturer webpage. b) Representative images of B6, WSB and NZO neurons in microfluidic chambers prior to injury, post-injury in the absence of ID-8, and post-injury in the presence of ID-8. c) Heatmap for the area occupied by axons in the chamber two (axonal side) before and after injury. d) Ratio of axonal growth compared to pre-injury in both control and treated neurons. Dunnet’s test was used for statistical analysis on three biological replicates. e) MDS plot showing the differences between cell lines over all three conditions. f) Barcode plots showing GO analysis of differential response to axonal injury between WSB and B6 in the absence or presence of ID-8. g) RTNA showing the top three regulons associated with the differences between WSB and B6 when axons are injured. h) Enrichment analysis and comparison of isoform switches between the differences in axonal recovery between B6 and WSB treated or not treated with ID8; i) The isoform switch in the *Ncoa4* gene. j) Axonal recovery experiment conducted on isolated primary cortical neurons from B6 and WSB postnatal day five mice. t-test for statistical analysis on three biological replicates.

Because of this divergent response to injury between B6 and WSB, we again undertook bulk RNA-seq analysis on these two cell lines to identify the underlying transcriptional differences. Interestingly, MDS analysis demonstrated vastly different responses between the two strains in the response to axotomy in the presence or absence of ID-8. As we saw from experiments in the maturation stage, ID-8 affected the transcriptome in both strains in control or axotomized cultures. However, when axons are severed, the B6 transcriptome showed limited change, irrespective of ID-8 treatment, whereas WSB transcriptome shifts markedly in response to axotomy in control and ID-8 treated cultures. The combination of ID-8 and axotomy displays an additive effect on the transcriptome in WSB, whereas in B6 this response is not observed, as axonal cut with or without ID-8 (“no drug”) elicits the same effect (Figure 9e).

When axons were cut, the two cell lines (B6 and WSB) expressed a core set of genes. These genes were *Map11*, *Ntsr1*, *Ccn5*, *Slc16a3* and, *Smtnl2* reflecting GO associated with GO terms cellular response to hypoxia and neuron projection, both being part of a successful neuronal reaction to injury (Figure 9f). Top regulons for the differences found between the cell lines when injured were *Ascl1*, Creb5, and *Olig2* (Figure 9g). These transcription factors are important for neuronal differentiation and myelination. *Ascl1* has been associated with a superior regenerative capacity in dorsal root ganglion neurons in CAST versus B6 *in vivo* ^57^. This differential expression pattern only emerged between the two strains after axonal injury.

After axons were cut and treated with ID-8, more genes were differentially expressed in WSB than in B6 (two-fold). Top common DEG genes in both cell lines were *Padi2*, *Calcr*, *Itgb4*, *Ifitm1*, *Npy*, *Tnfrsf11b*, *C1ra* and *Pcolce*; all of them are associated with axonal growth and guidance. In WSB-derived ID-8 treated neurons, gene sets associated with cell adhesion and central nervous system myelination were upregulated compared to the B6 counterpart (Figure 9h) and were confirmed by the number of hits per GO. This profile can explain a more robust axonal growth after axonal damage in WSB neurons. Finally, RTNA showed that two of the top three regulons associated with the differences in recovery after axonal growth, *Olig2* and *Ascl1,* were similar to those identified in the axonal damage response in the absence of ID-8. These results suggest that *Olig2*, *Creb5*, and *Ascl1* are part of the core regulatory network for a fast axonal growth after axonal damage when B6 and WSB are compared. Isoform switched analysis unveiled that *Ncoa4* was not only downregulated but also switched in isoforms was found after axotomy (Figure 9i). *Ncoa4* was recently associated with ferritinophagy and neurodegeneration ^58^.

A comparison of axonal regrowth of primary cortical neurons from P5 newborn B6 or WSB mice confirmed that the differences in growth post-injury seen in mESC derived neurons reflected the regrowth capacity of neurons isolated directly from the postnatal brain (Figure 9j).

Using all of the RNA-seq data, we analyzed the effect of ID-8 treatment independently of the cell strain, and also compared across the two strains independently of the treatment, throughout all stages analyzed. Analyzing effects of ID-8 in a strain independent fashion, we found only 15 DEG genes, but the cross-strain comparison independent of treatment found 261 DEG (Fig. S7a). Similar to the gene expression differences, differential use of transcript variants were found between strains, and our data indicated that in general, the consequences of isoform switches in B6 resulted in shorter transcripts (Fig. S7b). It is worth pointing out that *Gapdh,* a housekeeping gene commonly used to normalize expression data, was differentially expressed between WSB and B6 regardless of treatment (Fig. S7c). This disparity could be associated with differences in metabolic demands and energetic substrates. The latter correlates with differences we observed in other ontologies like mitochondrial respiratory chain complex II assembly and monocarboxylic acid metabolic process (data not shown). A gene set more enriched in B6 independently of treatment was the regulation of apoptotic process involved in development, whereas that a cerebral cortex development gene set was differentially affected by ID-8, B6 being more severely affected (Fig. S7d). Gene sets enriched in WSB relative to B6 in the presence of ID-8 included genes involved in cortex development, forebrain development, and cell proliferation in forebrain, all in accordance with differential effects on NPC proliferation in the two strains.

### Heterozygous loss of Dyrk1a is lethal on the B6 background but not on 129 or A/J

Finally, we sought to establish the relationship between the *in vitro* susceptibility of B6 to DYRK1A inhibition and the *in vivo* effects of *Dyrk1a* heterozygosity. We generated a null allele of *Dyrk1a* by crossing female mice carrying the germline CMV-Cre to male *Dyrk1a* mice carrying the conditional allele of *Dyrk1a* (floxed *Dyrk1a*) (Figure 10). When both strains were on the B6 background, we did not recover any live pups heterozygous for *Dyrk1a* deletion (*Dyrk1*a+/-) from this cross (0/93). We were able to obtain several mosaic offspring for the floxed and deleted alleles, but crossing male mosaic mice to wild-type B6 mice did not produce any live *Dyrk1a*+/-pups (0/46). Given that the prototype mouse model of *Dyrk1a* haploinsufficiency was maintained on a mixed B6/129 background ^59^, and our finding that B6 neural progenitors were uniquely sensitive to DYRK1A inhibition, we reasoned that mice carrying the deleted allele may survive on a different background. To test this idea, we crossed male mosaic mice to 129S1 female mice and obtained live *Dyrk1a*+/-mice on a mixed 129/B6 background. We then backcrossed *Dyrk1a*+/-mice to 129S1 for six generations and found that heterozygote animals were born at Mendelian ratios. We also mated B6 *Dyrk1* mosaic to AJ females, and live *Dyrk1a* heterozygous pups were born. These results indicate that heterozygous deletion of *Dyrk1a* causes early lethality on the B6 background but does not affect survival on the 129S1 or AJ background.

**Figure 10.**
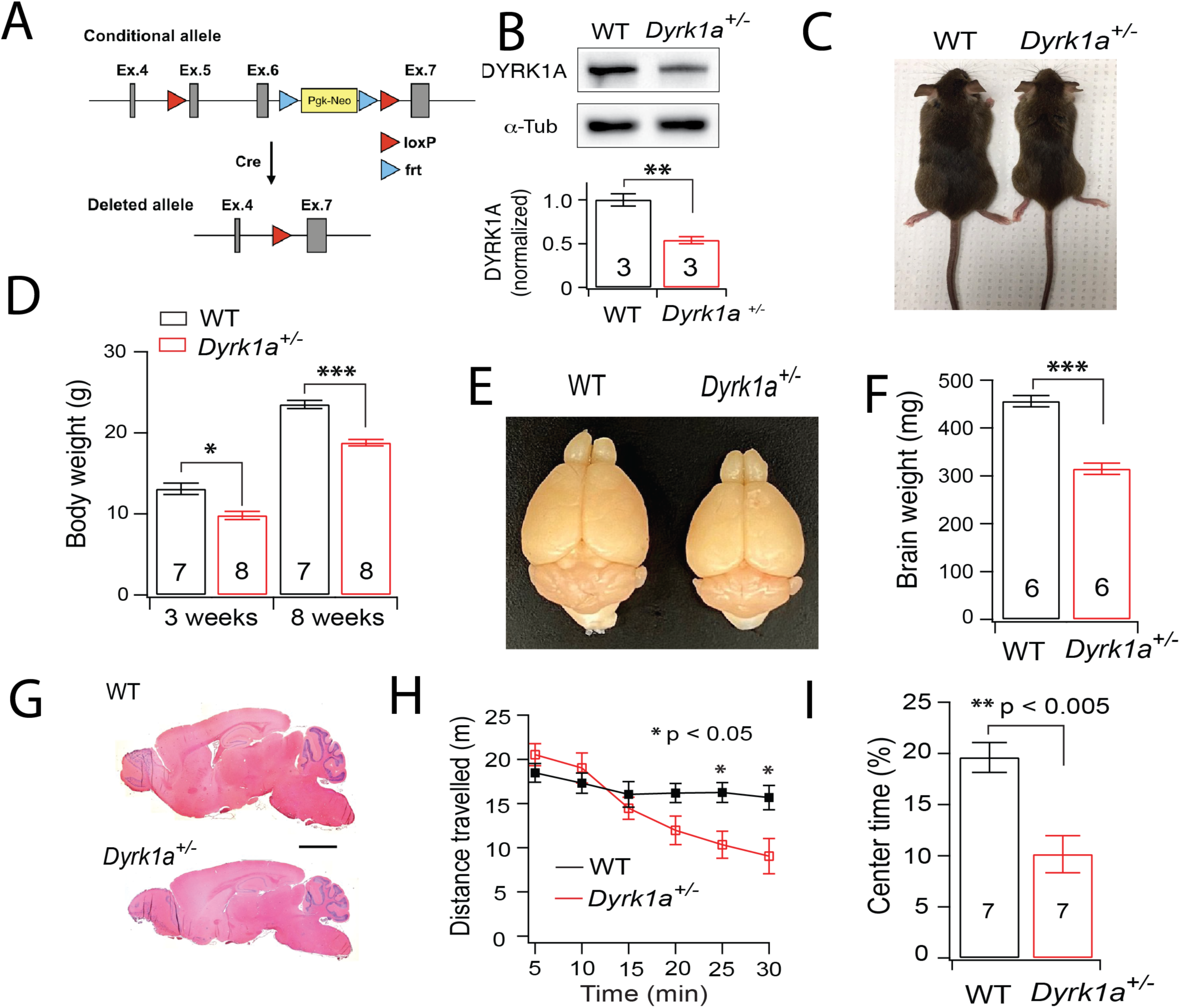
Heterozygous deletion of Dyrk1in mice confirms straind dependent sensitivity observed in vitro. a) Generation of the null allele of *Dyrk1a* using a conditional *Dyrk1a* allele and constitutively expressed *Cre*. Exons 5 and 6 of Dyrk1a, which encode part of the kinase domain, are flanked by loxP sites. The germline CMV Cre driver removes exons 5 and 6 and the Pgk-Neo selection cassette and causes a frameshift. b-i) Phenotype of *Dyrk1a* heterozygotes in B6/129 background. b) Western blot of cortical tissues from WT and Dyrk1a^+/-^ mice. α-Tub, α-tubulin. c) A Dyrk1a^+/-^ mouse and a WT littermate at 12 weeks of age. d) Body weight of male mutant and control mice at 3 and 12 weeks of age. e) Brains of a control and mutant at 12 weeks of age. f) Brain weights of mutant and control mice at 12 weeks of age. g) Sagittal brain sections from a WT and *Dyrk1a^+/-^* mouse at 3 months of age. Scale bar, 2 mm. h) Distance travelled by *Dyrk1a^+/-^* (n = 7) and WT (n = 7) mice per five minutes in an open field apparatus. i) Percentage of time spent at the center of the arena. * p < 0.05, ** p < 0.01, *** p < 0.001.

We further examined the phenotype of *Dyrk1a*^+/-^ mice on the 129B6F1 background. Western blot analyses showed that the level of DYRK1A protein was reduced by half in *Dyrk1a*^+/-^ mice (Figure 10b). Mutant mice appeared smaller and weighed 25% less than WT littermates during postnatal development and as young adults (Figure 10c,d). Likewise, the brains of *Dyrk1a*^+/-^ mice appeared smaller and weighed 30% less than those of control littermates (Figure 10e,f), although the brain to body ratio was not significantly changed. Brain histology indicated that the reduction in brain size was not uniform across the brain, with the mid and hindbrain regions showing larger reductions than the cerebral cortex (Figure 10g). These results corroborate previous findings that heterozygous deletion of *Dyrk1a* in the mouse causes growth retardation and microcephaly, two key clinical features associated with *DYRK1A* mutations in humans. *Dyrk1a^+/-^* mice on the 129B6F1 background also showed abnormal behaviors. In the open field apparatus, *Dyrk1a*^+/-^ mice were hypoactive compared with WT littermates (Figure 10h); they also spent less time in the center of the arena than WT littermates (Figure 10i), suggesting higher levels of anxiety in the mutant mice. Together, these findings confirm that heterozygous deletion of *Dyrk1a* on an appropriate genetic background recapitulates key features of human disease phenotypes, and that the B6 genetic background confers extreme sensitivity to the haploinsufficiency syndrome in the mouse.

## Discussion

To facilitate genetic analysis *in vitro* of neural development across a diverse range of mouse genetic backgrounds, we developed a new protocol to differentiate pluripotent stem cells into neurons from genetically diverse mESC and hPSC. We demonstrated the applicability of this model in analyzing the effects of chemical knockdown or genetic ablation of the activity of a key neurodevelopmental gene throughout neurogenesis and neural maturation and repair, and we were able to demonstrate how the knockdown phenotype varies dramatically depending on the genetic background.

The studies described here address three key questions in precision disease modeling. The first has to do with bridging the gap between *in vitro* studies and disease models in whole organisms, and asks, can disease modeling in genetically diverse mouse embryonic stem cells tell us what mouse strains are most representative of human phenotypes, and thus guide the choice of strain for whole organism studies? The second, a more general question of widespread significance, asks can any stem cell-based platform yield reliable information about the sensitivity or resilience of a particular individual genotype to pathogenic mutations? And the last question asks, does comparisons of *in vitro* phenotypes from stem cell models in sensitive and resilient strains yield reliable information about molecular pathways that might be critical to response to disease mutations in vivo?

Our studies have revealed that at most stages of this study, from embryonic stem cell renewal to neural specification, neurogenesis, and neural maturation and repair, B6-derived cultures were responsive to DYRK1A inhibition or loss while most other strains proved less sensitive. It is notable that C57BL/6J mice or hybrids thereof have been used successfully in a number of studies to model the effects of DYRK1A haploinsufficiency in human. Thus, on a C57BL/6J, 129S2/SvHsD mixed background, *Dyrk1a*^-/-^ mice show developmental delay and reduced brain size with greatly reduced neuronal content and die at mid-gestation^59^. Heterozygotes on this background have fewer intermediate neural progenitors in the subventricular zone at E11.5-13.5 owing to proliferative defects^60^. In the adult animal, cortical pyramidal neurons are smaller, and show less branching and fewer spines^61^, and neural progenitor maintenance in the subventricular zone is impaired^60^. These same animals show decreased cognitive abilities^62^. *Dyrk1a^-/+^* mice on a pure C57BL/6J background also show reduced brain size and cognitive defects^63^. Conditional knockout of *Dyrk1a* in the cortex of C57BL/6J mice was lethal in homozygotes; homozygotes and heterozygotes showed microcephaly related to reduced proliferation of neural progenitors in the immediate postnatal period, and a decreased complexity and reduced branching of cortical neurons^49^. We could not generate any heterozygous mouse on a pure C57BL/6J background, and only when it was crossed with less susceptible strains could we obtain heterozygous mice that showed similar effects to those described by other authors. Levy et al.^49^ used a cortex-specific KO approach which spared the remaining brain and body of the deleterious effects of the KO, making this intervention potentially less lethal. Raveau et al.^63^ used a strategy that was not tissue-specific, but the target region for the frameshift was upstream of the NLS and in exon 3, whereas in our case it was downstream of the NLS and the kinase domain was targeted.

It is possible that in our model, the remnant N-terminal peptide can cause a dominant-negative effect generating a more severe phenotype, or that, in the study of Raveau et al., exon skipping resulted in a protein with partial activity^63^. Interestingly, the observation that *Dyrk1a^-/+^* had smaller midbrain and hindbrain in our *in vivo* study correlates with aspects of our findings *in vitro*. We found that B6-derived NPCs treated with ID-8 had a decreased expression of *Hoxa4* and *Hoxd3*, homeotic genes associated with hindbrain development. Altogether, our results indicate an impaired development of the hindbrain at an early stage of neural differentiation unveiled by our *in vitro* analysis that otherwise would have been difficult to pinpoint.

Our previous work showed that neural specification of human pluripotent stem cells was blocked in *DYRK1A* knockdown or in ID-8 treated cells^35^. Here, we showed that the proliferative capacity of neural progenitors derived from *DYRK1A* +/-hPSC was reduced relative to controls. We observed both phenotypes in mESC or mESC-derived neural progenitors from B6 mice. These findings as well as the data discussed above show that our *in vitro* results correctly identify the B6 mouse as a sensitive model for the effects of *DYRK1A* haploinsufficiency in the human. By contrast, other strains were unresponsive to the developmental effects of DYRK1A inhibition. We also corroborated the effects of chemical DYRK1A inhibition by conducting knockout experiments demonstrating that the deleterious effect of DYRK1A inhibition is strain dependent. Although *DYRK1A* is subject to strong constraint in the human, and most heterozygotes show impairment of neural development, the severity of clinical phenotypes in individuals hemizygous for loss of the gene varies quite considerably^24,64^. Further *in vitro* analysis of mESC derived from recombinant inbred strains representing sensitive and resistant genetic backgrounds (B6 and 129 respectively) could identify key modifier genes, and potentially target pathways for therapeutic intervention during the early life of affected individuals. One such target is the BDNF signaling pathway, which we and others identify as a critical downstream modulator of *Dyrk1a* haploinsufficiency.

We also examined the effects of DYRK1A inhibition on axonal regrowth across the eight strains in our study. While repair and plasticity in the adult CNS often encompass some features of neurodevelopmental pathways, development and repair in the CNS are not driven by identical processes^65^. Our observations showed that the extent of axonal regrowth after injury in neurons derived from mESC does not directly relate to the extent of growth before injury across the mESC lines examined, suggesting that indeed axon growth during development and repair are different even in relatively immature neurons. There were large differences in the ability of axons derived from mESC of different strains to regrow in this assay. Importantly, the robust regrowth of neurons derived from the CAST strain contrasts strikingly with the poor regrowth of neurons derived from B6 mESC. Our observation of poor axonal regrowth in primary cultures of cortical neurons from P5 B6 or WSB mice confirmed the results obtained on neurons derived from mESC. In a number of studies *in vivo*, B6 mice have been shown to display deficient axonal regeneration relative to 129 or CAST ^66–71^, deficiencies that have been attributed either to variation in the inherent capacity for axonal repair or differences in the extent and nature of inflammatory responses in the various strains.

Comparison of gene expression in B6 and WSB strains that were sensitive or relatively resistant to effects of DYRK1A inhibition across different developmental stages revealed the power of the mESC model to identify novel critical molecular pathways and networks that are altered by genetic deletion of *Dyrk1A in vivo*. At the stage of neurogenesis, analysis of DEG highlighted the LIF pathway, which is a major regulator of neural progenitor proliferation *in vivo* not previously associated with Dyrk1a haploinsufficiency. RTNA found that networks related to *Foxb1*, a gene critical in the regulation of neural progenitor proliferation, were differentially active in the two strains. RTNA also identified two HOX genes involved in hindbrain development as targets of DYRK1A loss of activity in the B6 sensitive strain, consistent with phenotype data from B6/129 hybrid animals. Analysis of the expression of splice isoforms showed global changes in the presence of ID-8, in line with the known interactions of DYRK1A with splicing factors, and highlight alterations in splicing as an important determinant of susceptibility to *Dyrk1a* haploinsufficiency. The shift in isoform usage of the BDNF receptor *Ntrk2* to a form encoding a protein lacking the kinase domain in the presence of ID-8 in B6 neural progenitors could explain the downregulation of the BDNF pathway activity observed in a proteomics and phosphoproteomics study of conditional knockout of *Dyrk1a* in the cortical neurons of C57BL/6J mice^49^. Our studies of axonal repair identified the *Ascl1* regulon as a major differential response to injury in both strains either with or without DYRK1A inhibition. In a model of peripheral neuron (dorsal root ganglion) repair, Lisi et al. ^57^ noted a differential upregulation of *Ascl1* in CAST (identified by these authors and by our study as a good regenerator compared to B6) relative to B6. These authors also used overexpression and knockdown studies to confirm a role for *Ascl1* in axon regrowth. In summary, our *in vitro* studies comparing gene expression in strains sensitive and resistant to DYRK1A inhibition corroborate *in vivo* data on key molecular pathways involved in neuronal development and repair and their perturbations in *Dyrk1a*-deficient animals.

In our comparison of gene expression in both strains across all stages, the finding that *Gapdh* was the highest DEG was surprising, because it has been widely used as a reference housekeeping gene, with some studies supporting such use^72^ and others reaching the opposite conclusion^73^. GAPDH induces apoptosis when translocated into the nuclei of both neural and non-neural cells, and it shows dynamic changes in neurodegenerative diseases, cancer, and hypoxia^74,75^. GAPDH acts a sensor of NO stress^76^, and participates in dendrite development^74,75,77^. The strain differences we observed in *Gapdh* expression could have broader implications for disease modeling in B6 mice.

The approach we describe here could be generalized to address a range of issues in modeling the genetics of disease. Screening approaches using shRNAi or sgRNA CRISPR methodologies are also applicable in this system. We readily applied high content screening and medium throughput microelectrode array methodologies in our work. The small panel of mESC used in this study incorporates wide genetic diversity, but we are also developing mESC or iPSC panels from Diversity Outbred mice and from recombinant inbred strains. These mESC genetic diversity panels have been used for quantitative trait loci analysis, and could be used for rapid mapping of modifiers that influence phenotypes of genetic variants implicated in neurodevelopmental disorders. Identification of modifier genes could yield new leads for drug discovery and early intervention in patients. We focused here on neural development, but this same approach could be applied to any lineage or tissue that can be generated from pluripotent stem cells. Facile comparison between phenotypes of human and mouse pluripotent stem cell models *in vitro* will aid in the identification of those mouse genotypes that confer sensitivity or resilience to disease variants, providing for more precise disease modeling *in vivo* and reducing the experimental use of animals.

Finally, most disease modeling studies using hPSC have been retrospective, replicating the known effects of disease mutations to varying degrees in a culture dish. The use of hPSC and mESC in parallel followed by *in vivo* studies in the mouse provides a means for prospective testing of the ability of *in vitro* screens to predict the effects of genetic variants *in vivo*. This will be an important adjunct to hPSC studies aimed at validating novel candidate disease variants emerging from genetic studies in humans. Such a strategy will add considerable power to functional genomics analysis of variants of unknown significance, or variants for which the role in pathogenesis of disease is undefined. For such variants, establishing a link between observations in cell culture models and precision modeling *in vivo* will be essential.

Limitations and future directions: We acknowledge that we did not investigate the gender effect in our study and all stem cells are male-derived. Although ASD has a higher rate of affected males, and *DYRK1A*mutation is one of the causes of ASD^78^ ; *DYRK1A* haploinsufficiency syndrome does not seem to have a gender bias^79^. Evaluations of sex-dependent effects can be performed using female-derived stem cells, which are availabel from our laboratory; nonetheless, this is out of the scope of this paper. Also, we found that although our protocol can efficiently differentiate genetically diverse stem cells ot mature neurons, electrical maturation is achieved at different time rates as proven by our MEA recordings. Recently Hergenreder et al. https://doi.org/10.1101/2022.06.02.494616 reported that a small molecule cocktail can accelerate neuronal maturation. Addition of these molecules might synchronize the maturation rate of NPCs from genetically diverse stem cells.

## Materials and Methods

### RESOURCE AVAILABILITY

#### LEAD CONTACT

Further information and requests for resources and reagents should be directed to and will be fulfilled by the lead contact, Martin Pera.

#### MATERIALS AVAILABILITY

Dyrk1a^-/-^ mESC strains are available from our laboratory.

#### DATA AND CODE AVAILABILITY

- Bulk RNA-seq data have been deposited at GEO and are publicly available as of the date of publication. Accession numbers are listed in the key resources table.
- This paper does not report original code.
- Any additional information required to reanalyze the data reported in this paper is available from the lead contact upon request

#### EXPERIMENTAL MODEL AND SUBJECT DETAILS

##### ANIMALS

Experiments were conducted in accordance with the guidelines of the Institutional

Animal Care and Use Committee (IACUC) of the Jackson Laboratory. For axonal injury/recovery experiments in postnatal neurons, postnatal mice (p3-p5) from C57BL/6J and WSB/EiJ strains were sacrificed by decapitation and cells prepared according to the method below (Preparation of p5 NPC for axonal damage experiments).

##### CELL LINES

mESC from the CC founders^80^ were used; however, PWD/PhJ was substituted for PWK/PhJ. All mESCs were kept and used under ten passages to avoid chromosomal rearrangements. All mESCs are male, germ competent, negative for mycoplasma, and retain a normal karyotype. mESC media comprised Knockout DMEM, 15% Fetal calf serum, 2 mM Glutamax, non-essential amino-acids, 50 units/ml Penicillin, 50 μg/ml Streptomycin, 55 μM beta-mercaptoethanol, and 1000 IU of Leukemia inhibitory factor. Cells were grown on gamma-irradiated mouse embryonic fibroblast feeder cells at 37°C and 5% CO2. mESCs were passed using Trypsin 0.25%

KOLF2.1J hiPSC was used for neural differentiation studies comparing WT and heterozygous knock-out cells. CRISPR-Cas9 was used to produce frameshift mutants in Exon 2 on one allele of the *DYRK1A* gene, generating Dyrk1a +/-mutants, as previously described^81^. Cells were maintained on vitronectin-coated dishes in Stemflex media (Stem Cell Technologies) and passed using ReLSR (Stem Cell Technologies). hPSCs were kept and used under ten passages to avoid chromosomal rearrangements. hPSCs used in this study were karyotyped for chromosomal abnormalities and mycoplasma tested.

#### METHOD DETAILS

##### Cortical neuron differentiation of mESC

mESC were cultured as described above. Upon reaching 80% confluency (day 0), cells were trypsinized and seeded on a culture dish for an hour to let feeders attach to the dish. Unattached mESC were recovered and counted. EB were formed by seeding 3E5cells on a well of a 6-well plate (ultra-low attachment) in mESC media (described above in “CELL LINES”) plus 10 μM SB431542 and 1 μM Dorsomorphin. The next day (day 1), the EB-containing media was transferred to adherence; previously, a well of a 6-well plate was coated with 100 μg/mL Poly-Lysine (PLL) for 2 hours and then treated with, 1 μg/mL laminin (Lm), 5 μg/mL fibronectin (Fn) for at least two more hours. After 24h (day 2) of seeding the EB on adherance, media was removed and switched to NMM (Gaspard et al., 2009) (Neurobasal and DMEM/F12 1:1 vol/vol) plus SB431542, Dorsomorphin, 2 μM I-BET151, and 5 μM retinoic acid. 24h later (day 3), media was removed, and fresh NMM plus SB431542 and Dorsomorphin were added. The next day (day 4) SB431542 and Dorsomorphin were withdrawn for B6 and WSB, while the other strains were maintained on SB431542 and Dorsomorphin for two more days (until day 6) after which the inhibitors were withdrawn. Thereafter, all cells were kept on basal media (NMM) with daily media changes until day eight, when cells were passed to the NPC expansion stage (day 8 to 11). Dissociation of cells into small clumps was done using Tryple Express and expanded at a 1:2 split ratio on PLL/Fn/Lm-coated dishes in NMM plus 20 ng/mL FGF2, 20 ng/mL EGF, and RevitaCell (Gibco). After 24 hours (day 9), Revitacell is removed but mitogens are daily added for two more days. At day 11, cells were dissociated with Tryple Express to yield a pure population of NPC. NPCs were frozen using Knockout Serum Replacement and 10% DMSO, or passed onto a newly coated plate for neuronal maturation using adult NMM media (formulated with Neurobasal A in place of Neurobasal medium), plus Revitacell, 10 ng/mL GDNF, 10 ng/mL BDNF, and 10 μM Forskolin. Next day (day 12) Revitacell was removed. Three days later (day 15), Forskolin was removed and 5 μM 5-fluorodeoxyuridine was added for one week (until day 22). After that, adult-NMM + GDNF and BDNF was changed twice per week.

##### GABAergic differentiation of mESC

The first four days of differentiation were carried out as described in the cortical differentiation protocol above. On day five, SB431542 and Dorsomorphin were removed, and fresh media was added. On day six, a sonic hedgehog agonist (SAG, 10 μM) was added with every media change and withdrawn after three days of neuronal differentiation. The rest of the protocol was exactly the same as the cortical differentiation.

##### Dopaminergic differentiation of mESC

This procedure was the same as described for GABAergic neurons until day six. On this day, SAG and FGF8 (100 ng/ml) were added for two days, at which point conversion to NPC had occurred. Cells were passed as previously described, with the addition of Revitacell, FGF2, non-essential amino acids, FGF8, SAG1, IWP2 (1 μM) and LY364947 (3 μM) during the expansion stage. After plating cells for maturation, Revitacell, 10 μM Forskolin, BDNF, GDNF, 50 μg/ml ascorbic acid, CHIR99021 (3μM) and LY364947 were added. Media was changed on the third day with the continued addition of BDNF and GDNF. The cortical neural protocol was followed after day four of neuronal differentiation.

##### MN differentiation of mESC

Differentiation was as described for cortical neurons until day seven. On day four, SAG and retinoic acid were added with every change media. On day seven, neurons emerged and cells were ready to be seeded as young MN. When passing the cells, Revitacell, Forskolin, BDNF, GDNF, SAG1 and RA were added. After three days, all the small molecules were withdrawn and BDNF and GDNF were retained. After this point, half of the media volume was changed two to three times per week

##### Cerebral organoid differentiation of mESC

The protocol for organoid formation was based on the cortical differentiation protocol above. Briefly, 2000 cells were seeded per well of a round bottom, ultra-low attachment, 96-well plate in mESC basal media with LIF, SB431542, and Dorsomorphin. Forty-eight hours later, the spent media was removed and replaced with NMM+ SB431542, Dorsomorphin, I-BET151 and retinoic acid. The next day, this medium was removed and NMM with SB431542 and Dorsomorphin was added. Forty-eight hours later, half the volume of the culture medium was replaced. At day six and eight, half of the media was replaced with basal media (NMM). On day ten, organoids were split evenly and 24 organoids were placed into 6 cm ultra-low attachment dishes. On day 16, organoids are embedded in undiluted Matrigel (Corning, 356234) in the same medium and kept on a shaker at 16 rpm for at least 21 days.

##### hPSC-derived NPC

The protocol was the same as the one used for mESC with a few modifications . IBet151 and RA were added at day one instead of day two (still 24h treatment). SB431542, and Dorsomorphin were added until day eight. From day eight to eleven, cells were kept in basal NMM media. At day 11, the differentiated NPCs were passed and expanded using bFGF and EGF for three days. Afterwards, cells were frozen or used for further experiments.

##### hPSC Neurosphere assay

Human NPC were seeded on a 96-well ultra-low attachment plate at various concentrations (2, 8, 16 and 32 thousand cells) for 15 passages in NMM + FGF2 and EGF. Media changes were performed every other day. Spheroids were imaged with a Leica Dmi1 microscope and their size calculated using ImageJ software. Spheroids were passed every 4 days using Accutase for 30min at 37°C.

##### EdU labelling of NPC

Cells were incubated for 2 hours with 5µM EdU. After 48 hours, cells were harvested and fixed with 4% paraformaldehyde at room temperature for 15 minutes. EdU incorporation was detected according to the Clik-iT EdU Alexa Fluor 594 flow cytometry assay kit protocol. Cells not labelled with EdU were used as negative control.

##### hiPSC-derived cortical neurons

The protocol was similar as the one used for mESC but with slight modifications. Cells were dissociated with Accutase. 5E5 cells were seeded on a well of a low-attachment 6-well plate in the presence of SB431542, Dorsomorphin, Ibet151, RA, and Y-27632. 24 hours later, EB were sedimented under unit gravity, the media was removed, and 4 ml mTeSR plus media supplemented with SD434512 and Dorsomorphin was added. EB were seeded in a PLL/Fn/Lm coated well of the same size and incubated for 24 hours, after which the medium was replaced with NMM + SB and Dorsomorphin. This medium was changed every day until day 11, at which point small molecules were withdrawn incubation with basal media continued for 3 more days. Cells were dissociated into clumps of cells and reseeded at a 1:2 split ratio on PLL/Fn/Lm coated dishes in the presence of bFGF and EGF for two or three days for NPC expansion with the addition of Rock inhibitor during the first day. NPC can be frozen as with the mESC, or cells plated for neuronal differentiation, but 5FDU is added for seven days.

##### Differentiation of hPSC into motor neurons

Cell were differentiated according to the hPSC to cortical neuron protocol with minor modifications. On day eight, retinoic acid and SAG were added, and cells were maintained in this medium until day 23. Every time cells were passed, 5 μg/ml each of Fibronectin and Laminin were supplemented in the media in addition to their usage in the plate coating; these were supplemented again twice per week to avoid cell detachment and until 10 days after withdrawal of mitogens.

##### Differentiation of hPSC into Cerebral organoids

Initial differentiation was similar to 2D cortical cultures, but adapted for organoids by culturing in suspension, embedding in Matrigel and growth under agitation. Four thousand iPSCs were seeded in 100 μl of media per well of a 96 well plate with all aforementioned molecules.The following day, cultures were switched to mTeSR plus with SB and Dorsomorphin. Next day this medium was replaced with IDM (DMEM-F12 glutamax and Neurobasal 1:1 vol/vol ratio, N2 supplement, 5μg/ml insulin, non-essential amino acids, Penicillin-streptomycin and B27 supplement without Vit A) with SB and Dorsomorphin. This medium was changed every other day until day 11. At this point the organoids were embedded in Matrigel. The cultures were incubated in 6 ml of media overnight, and on the following day, transferred to to an orbital shaker with an orbit of 1.9 cm (0.75 inch) at 69 rpm. On day 18, half the media volume was replaced with fresh media. On day 20, Matrigel was removed and the medium switched to NMM. From then on, the media was changed two to three times a week. On day 30 cultures were switched to A-NMM adult with media changes two to three times per week.

##### Differentiation of hPSC into Dopaminergic organoids

Adaptation of the cortical organoid differentiation protocol was done as follow. SAG was added from day 4 until day 11 when it was withdrawn. FGF8 was added from day 6 until day 11 when it was withdrawn. LY36947 and IWP2 were added at day eight until day 11 and 15 respectively.CHIR99021 and Ascorbic Acid were added on day 11 until day 15 and day 30 respectively.

##### Calcium spike assay

Organoids were assayed using the FLIPR Calcium 6 assay kit (Molecular Devices). Briefly, organoids were incubated for two hours in half FLIPR Calcium 6 reconstituted reagent and half NMM media. Afterwards, this medium was removed and fresh media was added. Organoids were immediately transferred to Opera plus microscope (Perkin Elmer) under controlled environmental conditions (37C and 5% CO_2_ in a humidified chamber). Images were taken every 0.7 seconds up to 400 frames. Calcium spikes were analyzed using the integrated Harmony software by recognizing individual cells and tracking fluorophore intensity over time.

##### Electrophysiological Recordings

NPC were seeded on a previously coated MEA plate and left to mature as described above. However, for the MEA plates, a 10 μl droplet containing 50,000 mouse NPC or 20,000 human NPCs was seeded at the center of each well of a 48-well plate (CitoView MEA 48, Axion biosystems). After cells have attached (ý7h), more media was added to properly cover the well. Recordings were obtained on day 7, 14, 21, 28, 35, and 42. Spontaneous activity was recorded on an Axion Maestro Pro at 37°C for 10 minutes using the AxiS navigator software. Recordings were analyzed using the built-in spike detector at 6 x STD.

##### Axonal Damage using mESC-derived neurons

NPC of each cell line were seeded on one of the two chambers of a microfluidic device (ENUVIO). Devices were pre-coated with PLL/Fn/Lm. Cells were left to mature for four days. Afterwards, in the contralateral chamber, axons were cut off by scratching the chamber and pipetting the media up and down several times. The axonal chamber was rinsed to removed debris and replenished with fresh media and Fn/Lm to enhance axonal attachment. Four days later, the axonal side was treated with Calcein AM, and images of the entire device were taken. The area covered by axons was analyzed using ImageJ.

##### Preparation of p5 NPC for axonal damage experiments

C57BL/6J (B6) and WSB/EiJ (WSB) postnatal day five animals were euthanized as described above. The hippocampus/dentate gyrus was dissected, mechanically dissociated, and seeded on PLL/Fn/Lm-coated dishes in NMM media plus EGF and FGF (20 μg/ml each) and expanded through one passage. At this point, expanded NPC were used for axonal damage experiments in the same fashion as mESC-derived NPCs described above.

##### Immunostaining

Cultures were washed with PBS and fixed with 4% paraformaldehyde 4% for 15 min and finally washed again with PBS three more times. Blocking and permeabilization were carried out using 5% BSA and 0.3% Triton-X for an hour at room temperature. Primary antibodies used were OTX2, Millipore AB9566-1 (1:1000); TUBB3, Biolegend 801202 (1:1000); Ki67, Santa Cruz SC-23900 (1:50); PAX6, Biolegend 862001 (1:1000); Brachyury, R &D systems AF2085 (1:200); Vglut, Abcam ab104898 (1:200); CHAT, Merck Millipore AB144P (1:200); Hb9, Thermo PA5-67195 (1:200); MCM2, Abcam ab95361 (1:500); DYRK1A, cell, signaling, (1:1000); ISL1, DSHB 40.3A4-S and 40.2D6-S both undiluted 1:1; SOX2, Millipore AB5603 (1:500); Eomes, Abcam ab23345 (1:1000); POU5F1, BD 611202 (1:500); GATA4, Santa Cruz SC-25310 (1:2000); NANOG, PeproTech 500-P236-50UG (1:500); SOX1, R & D AF3369-SP (1:500); AFP, Dako A0008 (1:500); GFAP, Dako Z0334 (1:1000); NeuroD1, Abcam ab60704 (1:2000); TH, Pel-Freez P41301 (1:1000); GAD 65/67, Millipore ABN904 (1:250); vgat, Synaptic systems 131011 (1:200); Reelin, Abcam ab78540 (1:200). Primary antibodies were incubated overnight, washed three times with PBS and secondary ALEXA antibodies (1:1000) against primary antibodies were incubated for two hours. DAPI was used for nuclear counterstaining. Cerebral organoids were cleared and stained using fructose-glycerol clearing method (Dekkers et al., 2019) and antibodies were used at twice the concentrations described above.

##### High content microscopy

Perkin Elmer cell carrier ultra 96-well plates were used for high content microscopy. Imaging was carried out on the Operetta CSL (Perkin Elmer) microscope, and image analysis was done using Harmony software.

#### RT-PCR

Samples were collected and RNA was prepared using Qiagen RNAeasy plus mini kit (Qiagen). Reverse transcription was carried out using High-Capacity RNA-to-cDNA™ Kit (ThermoFisher). Real-time PCR was conducted using Quantstudio 7 Flex on a 2-step cycle run using Taqman Probes (Oct4, # 4331182; Nanog, #4331182; Otx2, #4331182; Pax6, #4331182; Tubb3, #4351372; Scl17a7, # 4331182; Gapdh, # 4453320).

##### Bulk RNA-seq

Samples were collected and RNA was prepared using Qiagen RNA column system. mRNA libraries were made at 150 x2 with at least 30 000 reads per sample.

##### Transcriptomic analysis

Low-quality bases (Q < 30) from the 3’ end of reads were removed and reads with more than 30% low-quality bases (Q < 30) overall were filtered out. Trimmed reads were aligned to the reference genome using bowtie2 ^82^, generating a bam file which was used by rsem for quantification providing expression counts for genes and isoforms separately. Quality metrics were obtained at the fastq and bam level. This includes metrics such as total number of reads, total number of high-quality reads and alignment rate. The RNA-seq data was filtered and normalized. MDS plots were made using EdgeR ^83^. Venn diagrams were made using systemPipeR ^84^ and the Limma ^85^ package was used for barcode plots and testing for multiple gene sets. Gene ontology analysis and plotting was completed using TopGO ^86^ and GO.db ^87^. Transcriptional network analysis was done using RTN ^88^. Circos plots were made using circlize ^89^. Isoform switching was analyzed using IsoformSwitchAnalyzeR package ^90^ and other tools to import data, conduct normalization, isoform switch testing, prediction of open reading frames, premature termination decay, coding, domaings signal peptides, protein disorder and consequences ^90–96^

##### Statistical analysis

Statistical details can be found within the figure legends. Computation and plots were done using Rstudio and GraphPad Prism.

##### SNP and transcript variant analysis

Dyrk1a sequences were pulled from the mouse genome project (https://www.mousegenomes.org/) and compared for number of SNPs and transcript variants per strains. Sequence variants were aligned using Clustal O multiple sequence alignment. Sequences downloaded from https://www.informatics.jax.org/mgv/

##### QTL analysis

Diversity outbred (DO) mouse embryonic stem cell (mESC) expression quantitative trait loci (eQTL) and protein QTL analysis are previously published (Skelly et al,. 2020 and Aydin et al., 2023)andavailableat (https://churchilllab.jax.org/qtlviewer/DO_mESC and https://doi.org/10.6084/m9.figshare.220128 <u>50</u>). Data for Dyrk1a was accessed through publicly available data sources. QTL mapping was performed using a linear-mixed model implemented in the ‘scan1’ function in R/qtl2 package with rank normal scores, sex of the cell lines as an additive covariate and the leave one chromosome out option for kinship correction (citations for R/qtl2 and DOQTL).

##### Western blot

Cells were lysed with NP40 cell lysis buffer supplemented with Protease Inhibitor Cocktail-P2714, Sigma. Cells were centrifuged for 15 min at 12,000g at 4°C. Supernatant was saved and kept frozen until further use. DC protein assay (Bio-Rad) was used to quantify protein. The cell lysate was placed in SDS sample buffer and run in a pre-casted gel at 100V. Proteins were transferred into nitrocellulose using the trans-Blot turbo (BIO-RAD). Primary antibodies are described above. Secondary antibodies were goat anti-Rb HRP (Abcam ab6721, 1:10,000) and horse anti-moouse (Vector Labs PI-2000-1). Detection was performed using SuperSignal Atto (Thermo) using a G box documentation system (Syngene).

##### Generation of *Dyrk1a* null allele

Mice carrying *Dyrk1a* null allele were generated by mating male mice homozygous of conditional *Dyrk1a* floxed allele (JAX Strain #027801) with female mice hemizygous of the germline CMV-Cre (JAX Strain #006054). Both strains were on a B6 background. Genotyping was performed by PCR on tail DNA. The PCR primers for Dyrk1a alleles are: forward 5’-CCT GGA GAA GAG GGC AAG-3’ (common), reverse 5’-GGC ATA ACTTGC ATA CAG TGG-3’ (wild type and floxed allele), and reverse 5’-ACT GTG TGA GGA GTC TTG ACA-3’ (null allele), which give products of 132, 232 and 295 bp for wild type, floxed, and null allele, respectively. The PCR primers for CMV-Cre are: forward 5’-GCA TTA CCG GTC GAT GCA ACG AGT GAT GAG-3’, reverse 5’-GAG TGA ACG AAC CTG GTC GAA ATC AGT GCG-3’, which gives a product of 410 bp for the transgene.

The cross between floxed *Dyrk1a* and CMV-Cre on B6 background did not produce any pups heterozygous of Dyrk1a null allele (0/93 pups), but a few live pups were mosaic for floxed and null allele. Crossing male mosaic mice to wild type B6 females did not produce any pups (0/46 pups). In contrast, crossing the same male mosaic mice to wild type 129S1 females produced live pups heterozygous of *Dyrk1a* null allele (*Dyrk1a+/-,* 15/81 pups). Further backcrossing to 129S1 produced *Dyrk1a*+/-mice at Mendelian ratios.

##### Phenotyping of *Dyrk1a* +/-mice

Mice on 129B6 F1 background were used for phenotyping. Western blots were performed with cortical tissues obtained from *Dyrk1a*+/- and wild type mice at 13 weeks of age, using standard procedures and antibodies against DYRK1A (Cell Signaling #8756, 1:1000) and ⍺-tubulin (ThermoFisher #A11126, 1:5000, loading control). Protein levels were quantified using ImageJ. For each sample, DYRK1A protein level was normalized to that of ⍺-tubulin.

Activity of mice was analyzed at 10 to 12 weeks of age with an open field apparatus where a single mouse was allowed to explore freely for 30 min and recorded by video. Activity was analyzed using idTracker (Perez-Escudero et al., Nature Methods, 2014). Distance travelled per 5 min and time spent in the center square were quantified.

For brain weight and morphology, mice were euthanized at 12 weeks of age and perfused with 4% paraformaldehyde. Whole brains were dissected and weighted. Brain sections were cut at 70 µm thickness and stained with H&E method.

##### *Dyrk1a* knockout mESC

C57Bl6/J mESC (line #49) were grown on irradiated mouse embryo fibroblasts in 2i/FCS/LIF media (Czechanski et al. 2014). Exon 3 of *Dyrk1a* (Dyrk1a-202, ENMUST00000119878) was deleted in ESC using CRISPR/Cas9. Briefly, oligos encoding guide RNAs targeting intron 2 and intron 3 of murine *Dyrk1a* were cloned into plasmid pX459 v2.0 (Addgene plasmid # 62988) and sequence verified. Plasmids encoding *Dyrk1a* sgRNAs were transfected into mouse ESC using Lipofectamine 3000 (Life/Thermo) at a ratio of 1 ug total plasmid per 200,000 cells. After 24 hours, cells were transferred to irradiated DR4 puromycin-resistant mouse embryo fibroblasts and transfected cells were selected for 40 hours with 1 ug/ml puromycin. Cells were then removed from puromycin selection and cultured for 1 week, then individual colonies were picked into 96 well plates. Clones were genotyped by PCR with primers Dyrk1a_For (5’-CATTCCATGCTGCTGGCCTTC-3’) and Dyrk1a_Rev: 5’-GCCAGCAGCATGGAATGAGAAC-3’) and analyzed by gel electrophoresis. Clones completely missing a wildtype band (1712 bp) were further expanded and re-genotyped for confirmation of loss of exon 3.

## Supporting information

CORTES ET AL. SUPPLEMENTAL FILES

## Acknowledgments

This work was supported by the Jackson Laboratory. DEC was the recipient of a JAX Scholar Award. AE was a participant in the JAX Summer Student Program. A National Institutes of Health grant to LGR (U42 OD010921) supported mESC derivation and expansion. We thank Dr. Catherine Kaczorowski for providing access to MEA equipment and for review of the manuscript, and Ms. Anne Czechanski for her advice on mESC culture. Ann Wells, Harshpreet Chandock, and Annat Haber (The Jackson Laboratory) advised on computational analyses. We thank Hong Liu for designing genotyping protocols and genotyping.

## Author Contributions

DEC: Conceptualization, Methodology, Validation, Formal Analysis, Investigation, Data Curation, Writing, Visualization, Supervision

ME: Investigation

ACK: Investigation

AE: Investigation

SCM: Formal Analysis, Visualization

SCA: Formal Analysis, Visualization

KC: Investigation

KMSO: Resources

LGR: Resources, Writing

MFP: Conceptualization, Methodology, Resources, Writing, Supervision. Project Administration, Funding Acquisition

## Supplementary Figure Legends

Fig. S1. Generation of a protocol to differentiate genetically diverse mESC into neural precursors. a) Diagram of the differentiation protocol to generate and expand NPC from mESC. b) Bright field images (left) and immunofluorescent images (right) of representative mESC differentiated into neural precursors in 8 days. NPC markers go up (Nestin), as well as forebrain markers (Pax6), pluripotency associated marker (Oct3/4) decreases completely while Sox2 remains expressed. c) Quantitative PCR for different pluripotency- and neuronal-associated markers at day 0, 2, 5 and 8 of differentiation of B6 strain mESC. Kruskal-Wallis and Dunn’s test were conducted. n = 5 in triplicate. * < p 0.5, ** < 0.1, *** < 0.001 compared to day zero. Data are represented as mean ± SEM.

Fig. S2. Heatmap of the percentage of cells positive for the analyzed markers throughout eight days of differentiation in the presence of ID-8.

Fig. S3. Genetically diverse hPSC differentiated into CO. a) Scheme of the differentiation protocol to generate hPSC-derived CO. b) Top panel, calcium imaging of hPSC cerebral organoids. Circles show two examples of cells with changes in calcium activity. Bottom panel, plot showing the activity of the two cell in the upper panel followed through time. c) Representative images of 33_ 1 hPSC-derived COs at 146 days old. D Quantification of the cortical layer markers found in the hPSC-derived COs. n = 4 biological replicates.

Fig. S4. Differentiation of hPSC into dopaminergic neurons. a) Scheme for differentiation of hPSC-derived dopaminergic organoids. b) Dopaminergic (TH) and midbrain (FOXA2) markers. .

Fig. S5. Genotypic and phenotypic analysis of Dyrk1a among the eight CC founder mESC lines. a) plots depicting the number of SNPs and transcriptional variants among CC founder mESC compared to the reference genome B6. b) eQTL for both mESC and differentiated NPC show a peak in the Dyrk1a locus representing a local eQTL that can drive differences in expression specially in PWK and CAST. c) pQTL shows no differences in protein expression among all strains analyzed. d, e) Western blot shows protein abundance among mESC strains but no statistical differences.

Fig. S6. a) MDS plot comparing B6- and WSB-derived neurons treated or not with ID-8. It shows the distance between the groups and similarities in gene expression from RNA-seq samples. b) Venn diagram of DE genes for each set of comparisons. Number in red represents downregulated genes, black indicates upregulated genes in B6 compared to WSB. c) Exemplary barcode plots for differentially regulated gene sets. Conditions compared are indicated on both sides of the barcodes. d) RTNA for the top TFs associated with differences in ID-8 treated WSB- and B6-derived neurons. e) Alternative splicing analysis showing the change in alternative transcript start site (ATSS), alternative transcript termination site (ATTS), and alternative donor and acceptor sites (A5, A3) when B6 and WSB neurons are treated with ID8. f) Isoform switch for Slit2 in B6-derived neurons when treated with ID-8.

Fig. S7. Comparative transcriptomic analysis across all datasets. a) Venn diagram showing the numbers of differentially expressed genes, strain-wise and treatment-wise; b) Consequences of switch analysis between the B6 and WSB strains; c) Relative expression of *Gapdh* in both strains. Two-tailed T test, n = 38, *** = p < 0.001; D) Bar code plots showing overall differences between the two strains in control and when treated with ID-8.

