## Supplementary material for "An *in vitro* neurogenetics platform for precision disease modeling in the mouse": CORTES ET AL. SUPPLEMENTAL FILES

**This PDF file includes:**

Figs. S1 to S7
Tables S1

**Other Supplementary Materials for this manuscript include the following:**
Movies S1

FIG. S1

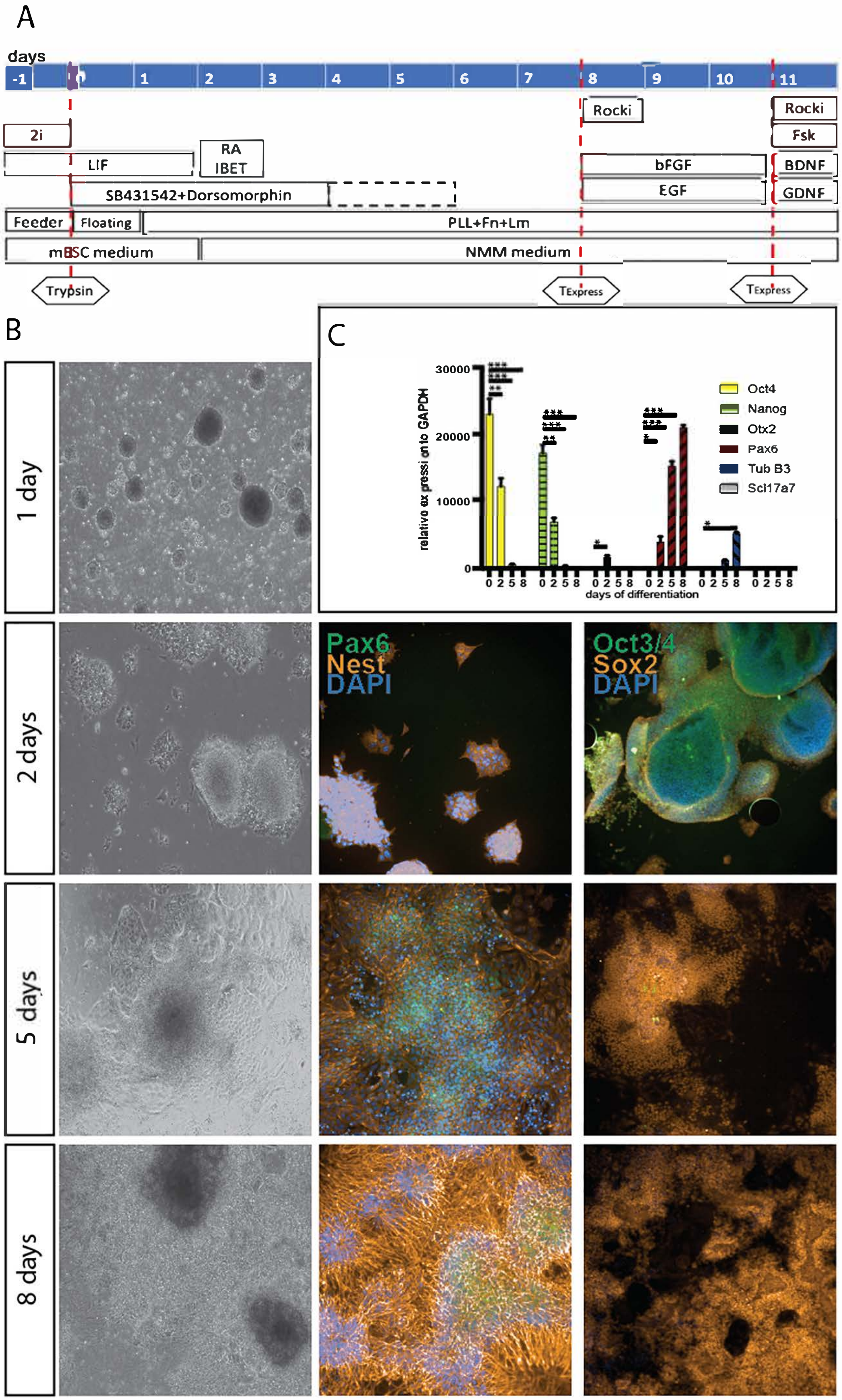


Fig. S1. Generation of a protocol to differentiate genetically diverse mESC into neural precursors. a) Diagram of the differentiation protocol to generate and expand NPC from mESC. b) Bright field images (left) and immunofluorescent images (right) of representative mESC differentiated into neural precursors in 8 days. NPC markers go up (Nestin), as well as forebrain markers (Pax6), pluripotency associated marker (Oct3/4) decreases completely while Sox2 remains expressed. c) Quantitative PCR for different pluripotency- and neuronal-associated markers at day 0, 2, 5 and 8 of differentiation of B6 strain mESC. Kruskal-Wallis and Dunn's test were conducted. n = 5 in triplicate. * < p 0.5, ** < 0.1, *** < 0.001 compared to day zero. Data are represented as mean ± SEM.

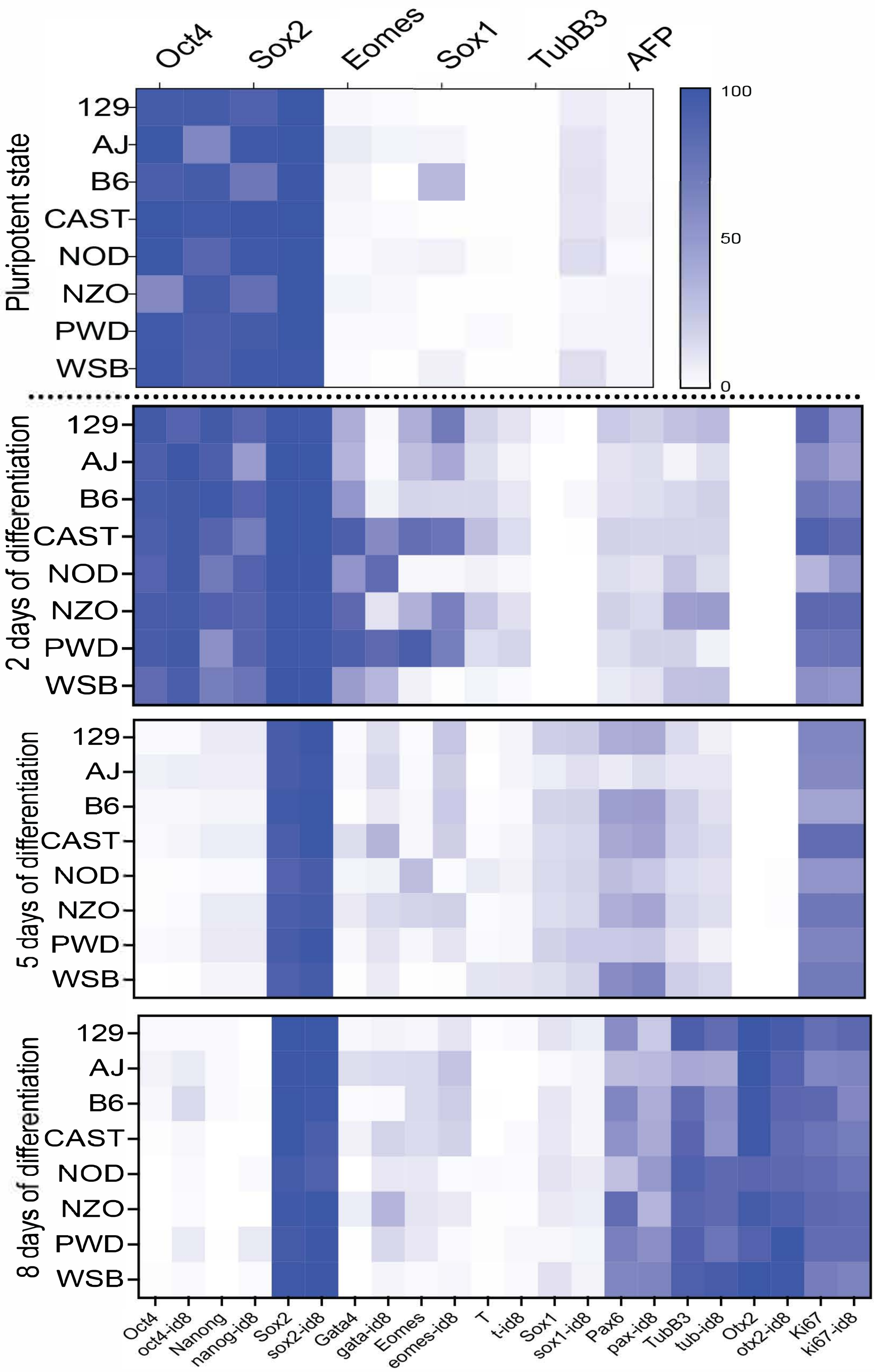


Fig. S2. Heatmap of the percentage of cells positive for the analyzed markers throughout eight days of differentiation in the presence of 1D-8.

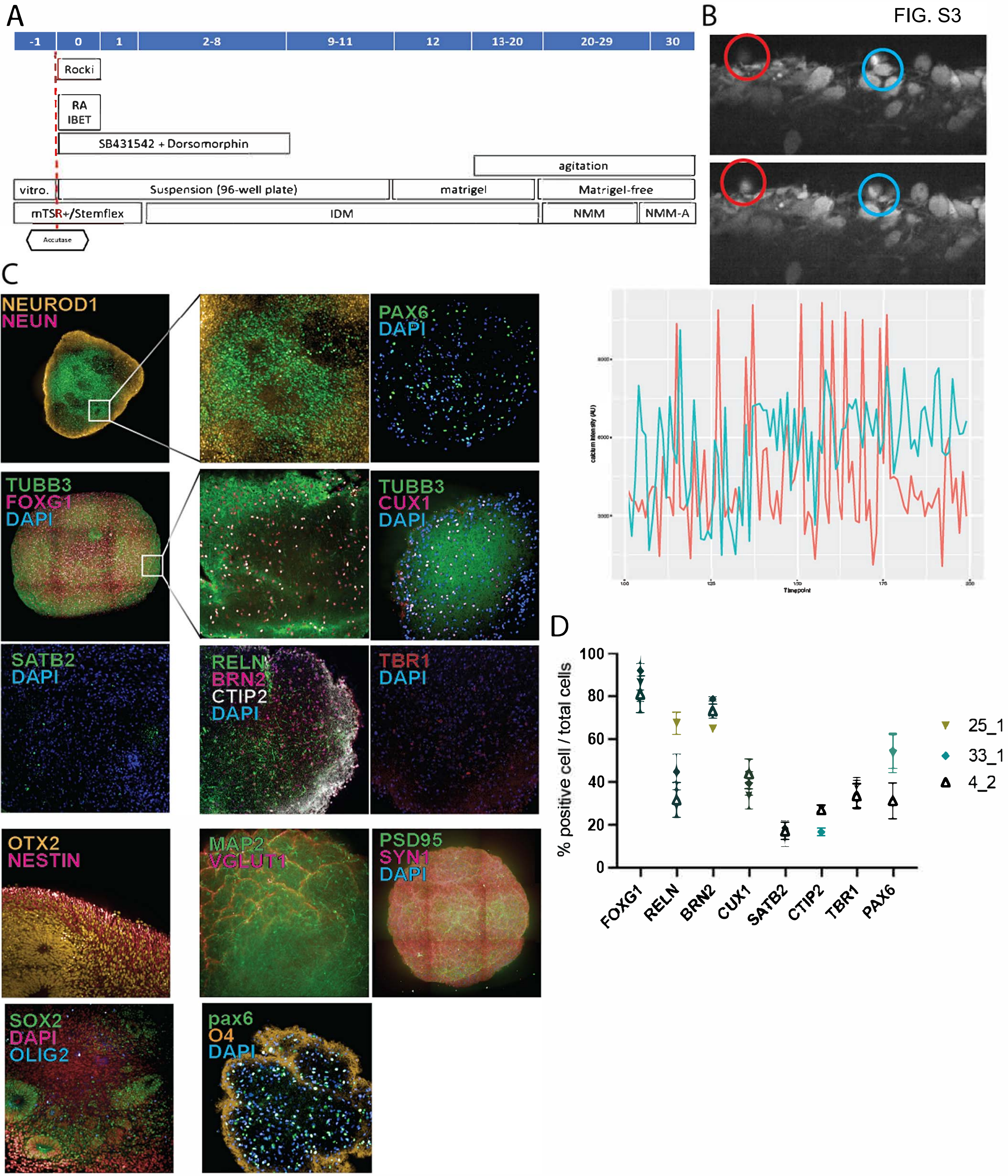


Fig. S3 Genetically diverse hPPSC differentiated into CO. A) Scheme of the differentiation protocol to generate hPSC-derived CO. B) Top panel, calcium imaging of hPSC cerebral organoids. Circles show two examples of cells with changes in calcium activity. Bottom panel, plot showing the activity of the two cell in the upper panel followed through time. C) Representative images of 33_1 hPSC-derived COs at 146 days old. D Quantification of the cortical layer markers found in the hPSC-derived COs. n = 4 biological replicates.

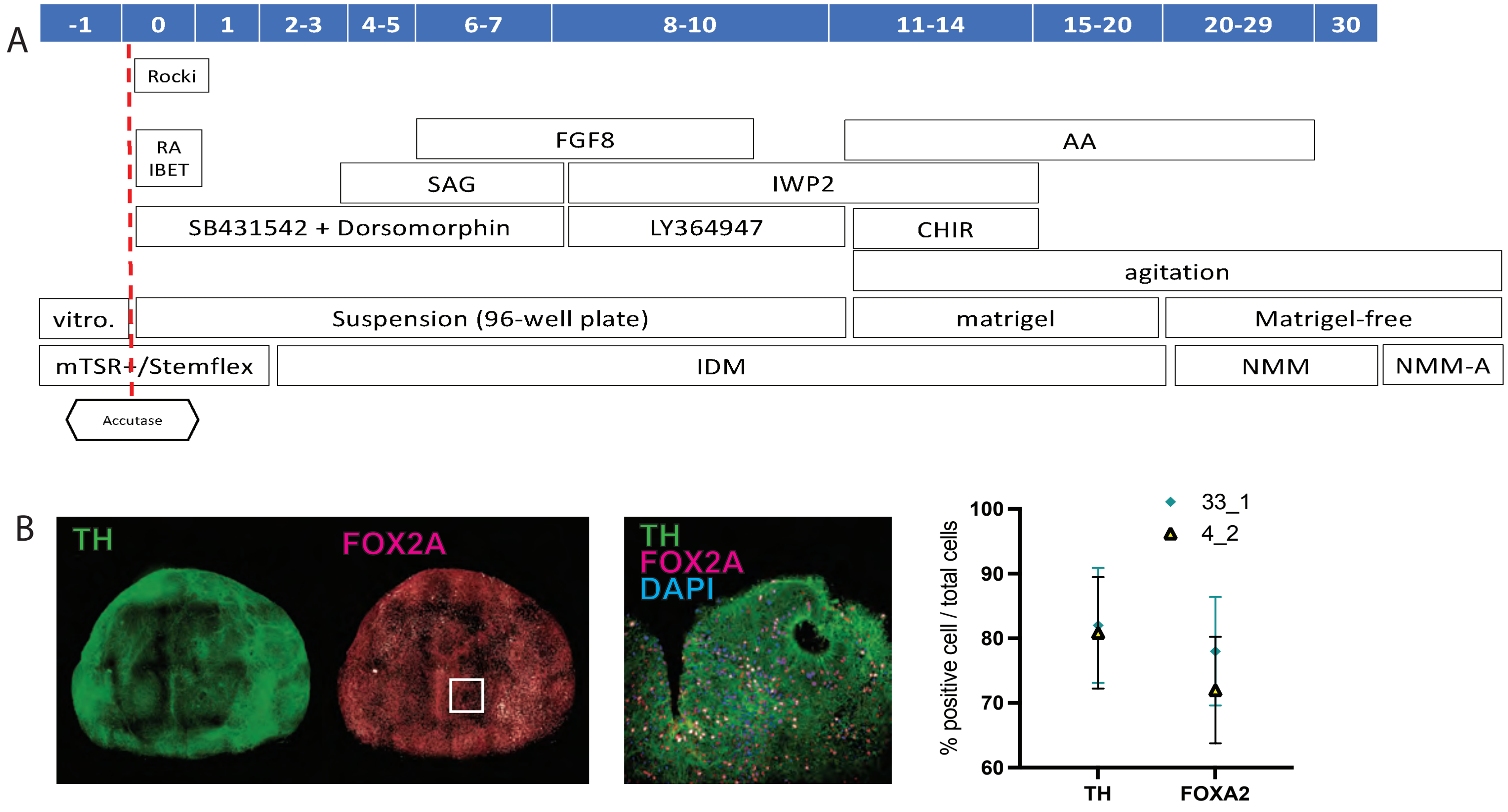


Fig. S4. Differentiation of hPSC into dopaminergic neurons. a) Scheme for differentiation of hPSC-derived dopaminergic organoids. b) Dopaminergic (TH) and midbrain (FOXA2) markers.

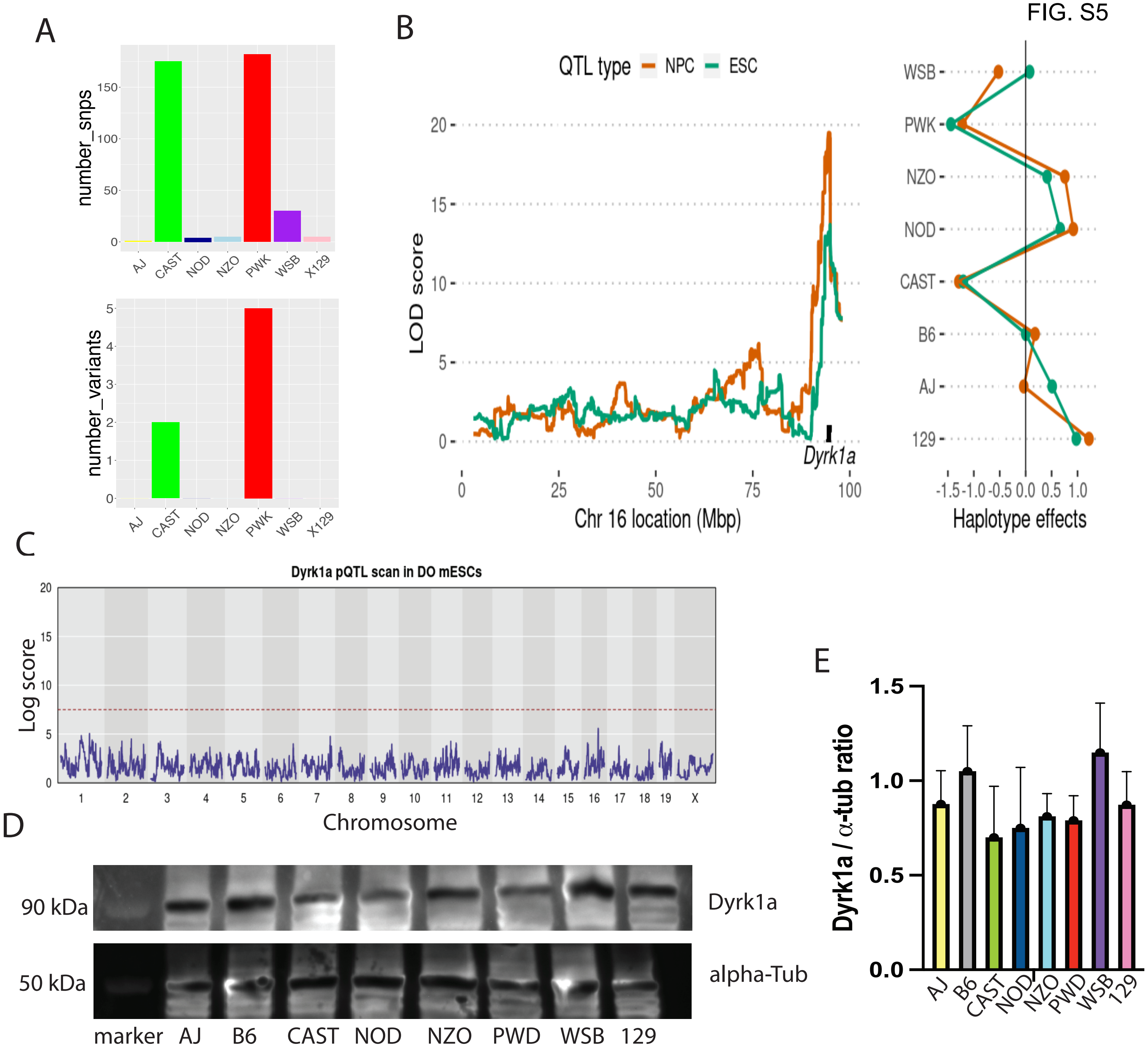


Fig. S5. Genotypic and phenotypic analysis of Dyrk1a among the eight CC founder mESC lines. a) plots depicting the number of SNPs and transcriptional variants among CC founder mESC compared to the reference genome B6. b) eQTL for both mESC and differentiated NPC show a peak in the Dyrk1a locus representing a local eQTL that can drive differences in expression specially in PWK and CAST. c) pQTL shows no differences in protein expression among all strains analyzed. d,e) Western blot shows protein abundance among mESC strains but no statistical differences.

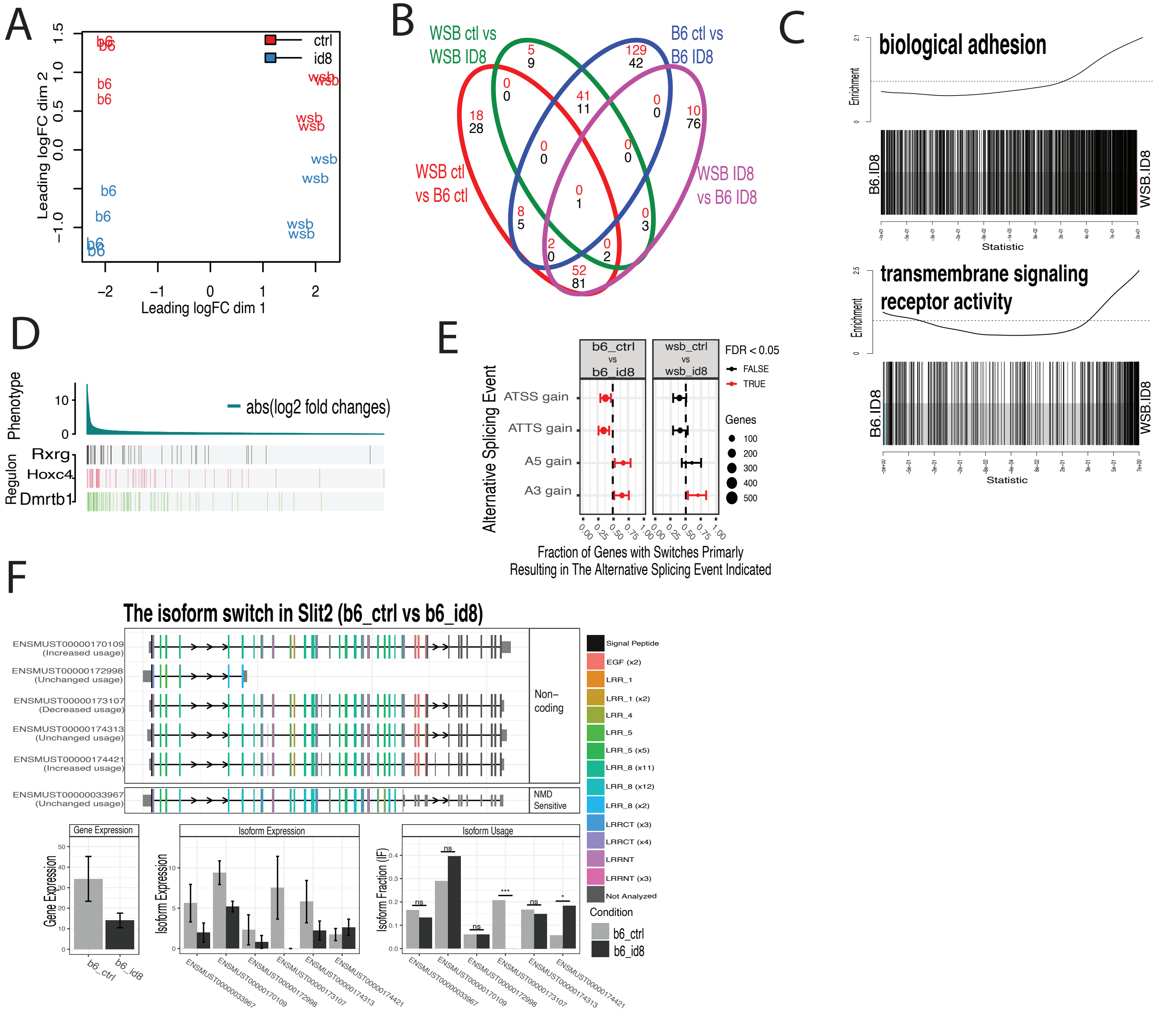


Fig. S6. a) MDS plot comparing B6- and WSB-derived neurons treated or not with ID-8. It shows the distance between the groups and similarities in gene expression from RNA-seq samples. b) Venn diagram of DE genes for each set of comparisons. Number in red represents downregulated genes, black indicates upregulated genes in B6 compared to WSB. c) Exemplary barcode plots for differentially regulated gene sets. Conditions compared are indicated on both sides of the barcodes. d) RTNA for the top TFs associated with differences in ID-8 treated WSB- and B6-derived neurons. e) Alternative splicing analysis showing the change in alternative transcript start site (ATSS), alternative transcript termination site (ATTS), and alternative donor and acceptor sites (A5, A3) when B6 and WSB neurons are treated with ID8. f) Isoform switch for Slit2 in B6-derived neurons when treated with ID-8.

FIG. S7

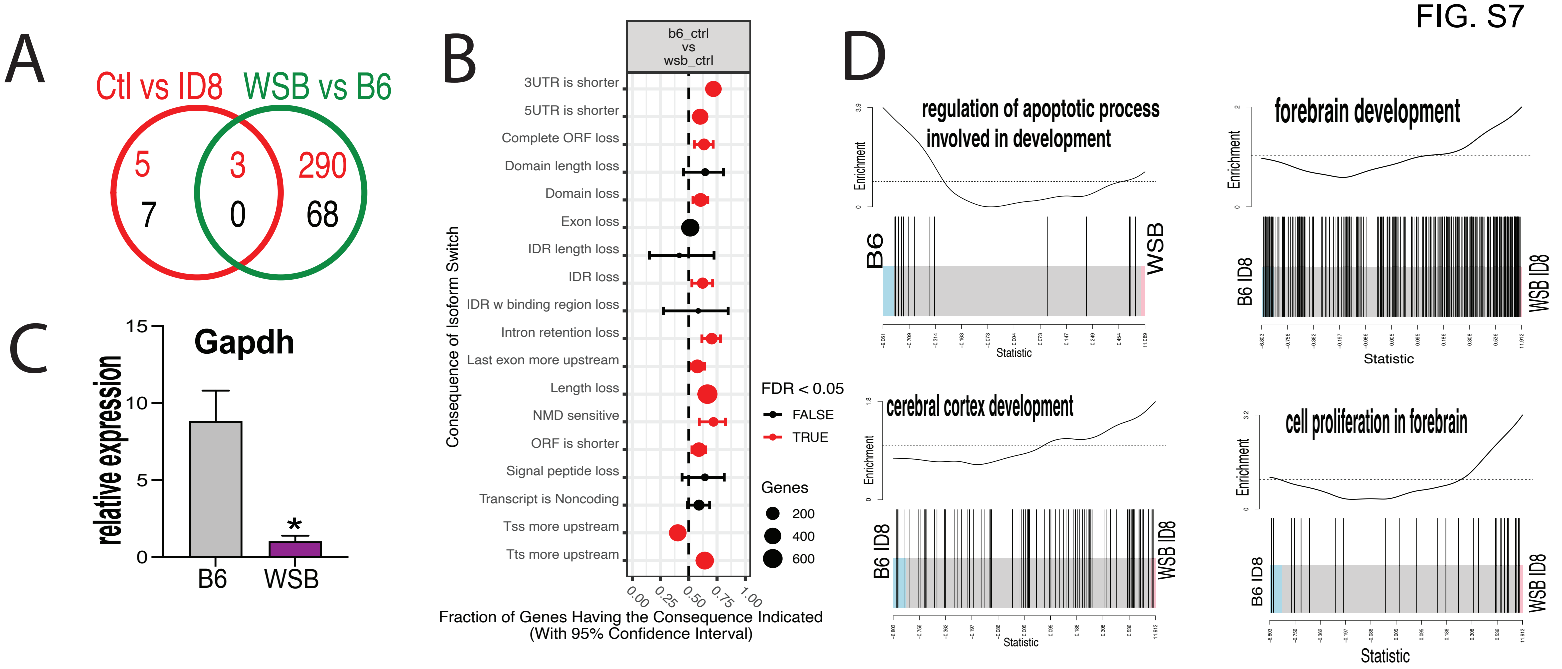


Fig. S7. Comparative transcriptomic analysis across all datasets. a) Venn diagram showing the numbers of differentially expressed genes, strain-wise and treatment-wise; b) Consequences of switch analysis between the B6 and WSB strains; c) Relative expression of Gapdh in both strains. Two-tailed T test, n = 38, *** = p < 0.001; D) Bar code plots showing overall differences between the two strains in control and when treated with ID-8.

```
CLUSTAL O(1.2.4) multiple sequence alignment


CAST/EiJ_MGP_CASTEiJ_P0045101
MHTGGETSACKPSSVRLAPSFSFHAAGLQMAAQMPHSHQYSDRRQPSISDQQVSALPYSD    60
129S1/SvImJ_MGP_129S1SvImJ_P0045278
MHTGGETSACKPSSVRLAPSFSFHAAGLQMAAQMPHSHQYSDRRQPSISDQQVSALPYSD    60
NZO/HlLtJ_MGP_NZOHlLtJ_P0045778
MHTGGETSACKPSSVRLAPSFSFHAAGLQMAAQMPHSHQYSDRRQPSISDQQVSALPYSD    60
NOD/ShiLtJ_MGP_NODShiLtJ_P0044977
MHTGGETSACKPSSVRLAPSFSFHAAGLQMAAQMPHSHQYSDRRQPSISDQQVSALPYSD    60
PWK/PhJ_MGP_PWKPhJ_P0044675
MHTGGETSACKPSSVRLAPSFSFHAAGLQMAAQMPHSHQYSDRRQPSISDQQVSALPYSD    60
A/J_MGP_AJ_P0045265
MHTGGETSACKPSSVRLAPSFSFHAAGLQMAAQMPHSHQYSDRRQPSISDQQVSALPYSD    60
WSB/EiJ_MGP_WSBEiJ_P0044346
MHTGGETSACKPSSVRLAPSFSFHAAGLQMAAQMPHSHQYSDRRQPSISDQQVSALPYSD    60
C57BL/6J_NP_031916.1
MHTGGETSACKPSSVRLAPSFSFHAAGLQMAAQMPHSHQYSDRRQPSISDQQVSALPYSD    60

************************************************************

CAST/EiJ_MGP_CASTEiJ_P0045101
QIQQPLTNQVMPDIVMLQRRMPQTFRDPATAPLRKLSVDLIKTYKHINEVYYAKKKRRHQ    120
129S1/SvImJ_MGP_129S1SvImJ_P0045278
QIQQPLTNQVMPDIVMLQRRMPQTFRDPATAPLRKLSVDLIKTYKHINEVYYAKKKRRHQ    120
NZO/HlLtJ_MGP_NZOHlLtJ_P0045778
QIQQPLTNQVMPDIVMLQRRMPQTFRDPATAPLRKLSVDLIKTYKHINEVYYAKKKRRHQ    120
NOD/ShiLtJ_MGP_NODShiLtJ_P0044977
QIQQPLTNQVMPDIVMLQRRMPQTFRDPATAPLRKLSVDLIKTYKHINEVYYAKKKRRHQ    120
PWK/PhJ_MGP_PWKPhJ_P0044675
QIQQPLTNQVMPDIVMLQRRMPQTFRDPATAPLRKLSVDLIKTYKHINEVYYAKKKRRHQ    120
A/J_MGP_AJ_P0045265
QIQQPLTNQVMPDIVMLQRRMPQTFRDPATAPLRKLSVDLIKTYKHINEVYYAKKKRRHQ    120
WSB/EiJ_MGP_WSBEiJ_P0044346
QIQQPLTNQVMPDIVMLQRRMPQTFRDPATAPLRKLSVDLIKTYKHINEVYYAKKKRRHQ    120
C57BL/6J_NP_031916.1
QIQQPLTNQVMPDIVMLQRRMPQTFRDPATAPLRKLSVDLIKTYKHINEVYYAKKKRRHQ    120

************************************************************

CAST/EiJ_MGP_CASTEiJ_P0045101
QGQGDDSSHKKERKVYNDGYDDDNYDYIVKNGEKWMDRYEIDSLIGKGSFGQVVKAYDRV    180
129S1/SvImJ_MGP_129S1SvImJ_P0045278
QGQGDDSSHKKERKVYNDGYDDDNYDYIVKNGEKWMDRYEIDSLIGKGSFGQVVKAYDRV    180
NZO/HlLtJ_MGP_NZOHlLtJ_P0045778
QGQGDDSSHKKERKVYNDGYDDDNYDYIVKNGEKWMDRYEIDSLIGKGSFGQVVKAYDRV    180
NOD/ShiLtJ_MGP_NODShiLtJ_P0044977
QGQGDDSSHKKERKVYNDGYDDDNYDYIVKNGEKWMDRYEIDSLIGKGSFGQVVKAYDRV    180
PWK/PhJ_MGP_PWKPhJ_P0044675
QGQGDDSSHKKERKVYNDGYDDDNYDYIVKNGEKWMDRYEIDSLIGKGSFGQVVKAYDRV    180
A/J_MGP_AJ_P0045265
QGQGDDSSHKKERKVYNDGYDDDNYDYIVKNGEKWMDRYEIDSLIGKGSFGQVVKAYDRV    180
WSB/EiJ_MGP_WSBEiJ_P0044346
QGQGDDSSHKKERKVYNDGYDDDNYDYIVKNGEKWMDRYEIDSLIGKGSFGQVVKAYDRV    180
C57BL/6J_NP_031916.1
QGQGDDSSHKKERKVYNDGYDDDNYDYIVKNGEKWMDRYEIDSLIGKGSFGQVVKAYDRV    180

************************************************************

CAST/EiJ_MGP_CASTEiJ_P0045101
EQEWVAIKIIKNKKAFLNQAQIEVRLLELMNKHDTEMKYYIVHLKRHFMFRNHLCLVFEM    240
129S1/SvImJ_MGP_129S1SvImJ_P0045278
EQEWVAIKIIKNKKAFLNQAQIEVRLLELMNKHDTEMKYYIVHLKRHFMFRNHLCLVFEM    240
NZO/HlLtJ_MGP_NZOHlLtJ_P0045778
EQEWVAIKIIKNKKAFLNQAQIEVRLLELMNKHDTEMKYYIVHLKRHFMFRNHLCLVFEM    240
NOD/ShiLtJ_MGP_NODShiLtJ_P0044977
EQEWVAIKIIKNKKAFLNQAQIEVRLLELMNKHDTEMKYYIVHLKRHFMFRNHLCLVFEM    240
PWK/PhJ_MGP_PWKPhJ_P0044675
EQEWVAIKIIKNKKAFLNQAQIEVRLLELMNKHDTEMKYYIVHLKRHFMFRNHLCLVFEM    240
A/J_MGP_AJ_P0045265
EQEWVAIKIIKNKKAFLNQAQIEVRLLELMNKHDTEMKYYIVHLKRHFMFRNHLCLVFEM    240
WSB/EiJ_MGP_WSBEiJ_P0044346
EQEWVAIKIIKNKKAFLNQAQIEVRLLELMNKHDTEMKYYIVHLKRHFMFRNHLCLVFEM    240
C57BL/6J_NP_031916.1
EQEWVAIKIIKNKKAFLNQAQIEVRLLELMNKHDTEMKYYIVHLKRHFMFRNHLCLVFEM    240

************************************************************

CAST/EiJ_MGP_CASTEiJ_P0045101
```

```
LSYNLYDLLRNTNFRGVSLNLTRKFAQQMCTALLFLATPELSIIHCDLKPENILLCNPKR    300
129S1/SvImJ_MGP_129S1SvImJ_P0045278
LSYNLYDLLRNTNFRGVSLNLTRKFAQQMCTALLFLATPELSIIHCDLKPENILLCNPKR    300
NZO/HlLtJ_MGP_NZOHlLtJ_P0045778
LSYNLYDLLRNTNFRGVSLNLTRKFAQQMCTALLFLATPELSIIHCDLKPENILLCNPKR    300
NOD/ShiLtJ_MGP_NODShiLtJ_P0044977
LSYNLYDLLRNTNFRGVSLNLTRKFAQQMCTALLFLATPELSIIHCDLKPENILLCNPKR    300
PWK/PhJ_MGP_PWKPhJ_P0044675
LSYNLYDLLRNTNFRGVSLNLTRKFAQQMCTALLFLATPELSIIHCDLKPENILLCNPKR    300
A/J_MGP_AJ_P0045265
LSYNLYDLLRNTNFRGVSLNLTRKFAQQMCTALLFLATPELSIIHCDLKPENILLCNPKR    300
WSB/EiJ_MGP_WSBEiJ_P0044346
LSYNLYDLLRNTNFRGVSLNLTRKFAQQMCTALLFLATPELSIIHCDLKPENILLCNPKR    300
C57BL/6J_NP_031916.1
LSYNLYDLLRNTNFRGVSLNLTRKFAQQMCTALLFLATPELSIIHCDLKPENILLCNPKR    300

************************************************************

CAST/EiJ_MGP_CASTEiJ_P0045101
SAIKIVDFGSSCQLGQRIYQYIQSRFYRSPEVLLGMPYDLAIDMWSLGCILVEMHTGEPL    360
129S1/SvImJ_MGP_129S1SvImJ_P0045278
SAIKIVDFGSSCQLGQRIYQYIQSRFYRSPEVLLGMPYDLAIDMWSLGCILVEMHTGEPL    360
NZO/HlLtJ_MGP_NZOHlLtJ_P0045778
SAIKIVDFGSSCQLGQRIYQYIQSRFYRSPEVLLGMPYDLAIDMWSLGCILVEMHTGEPL    360
NOD/ShiLtJ_MGP_NODShiLtJ_P0044977
SAIKIVDFGSSCQLGQRIYQYIQSRFYRSPEVLLGMPYDLAIDMWSLGCILVEMHTGEPL    360
PWK/PhJ_MGP_PWKPhJ_P0044675
SAIKIVDFGSSCQLGQRIYQYIQSRFYRSPEVLLGMPYDLAIDMWSLGCILVEMHTGEPL    360
A/J_MGP_AJ_P0045265
SAIKIVDFGSSCQLGQRIYQYIQSRFYRSPEVLLGMPYDLAIDMWSLGCILVEMHTGEPL    360
WSB/EiJ_MGP_WSBEiJ_P0044346
SAIKIVDFGSSCQLGQRIYQYIQSRFYRSPEVLLGMPYDLAIDMWSLGCILVEMHTGEPL    360
C57BL/6J_NP_031916.1
SAIKIVDFGSSCQLGQRIYQYIQSRFYRSPEVLLGMPYDLAIDMWSLGCILVEMHTGEPL    360

************************************************************

CAST/EiJ_MGP_CASTEiJ_P0045101
FSGANEVDQMNKIVEVLGIPPAHILDQAPKARKFFEKLPDGTWSLKKTKDGKREYKPPGT    420
129S1/SvImJ_MGP_129S1SvImJ_P0045278
FSGANEVDQMNKIVEVLGIPPAHILDQAPKARKFFEKLPDGTWSLKKTKDGKREYKPPGT    420
NZO/HlLtJ_MGP_NZOHlLtJ_P0045778
FSGANEVDQMNKIVEVLGIPPAHILDQAPKARKFFEKLPDGTWSLKKTKDGKREYKPPGT    420
NOD/ShiLtJ_MGP_NODShiLtJ_P0044977
FSGANEVDQMNKIVEVLGIPPAHILDQAPKARKFFEKLPDGTWSLKKTKDGKREYKPPGT    420
PWK/PhJ_MGP_PWKPhJ_P0044675
FSGANEVDQMNKIVEVLGIPPAHILDQAPKARKFFEKLPDGTWSLKKTKDGKREYKPPGT    420
A/J_MGP_AJ_P0045265
FSGANEVDQMNKIVEVLGIPPAHILDQAPKARKFFEKLPDGTWSLKKTKDGKREYKPPGT    420
WSB/EiJ_MGP_WSBEiJ_P0044346
FSGANEVDQMNKIVEVLGIPPAHILDQAPKARKFFEKLPDGTWSLKKTKDGKREYKPPGT    420
C57BL/6J_NP_031916.1
FSGANEVDQMNKIVEVLGIPPAHILDQAPKARKFFEKLPDGTWSLKKTKDGKREYKPPGT    420

************************************************************

CAST/EiJ_MGP_CASTEiJ_P0045101
RKLHNILGVETGGPGGRRAGESGHTVADYLKFKDLILRMLDYDPKTRIQPYYALQHSFFK    480
129S1/SvImJ_MGP_129S1SvImJ_P0045278
RKLHNILGVETGGPGGRRAGESGHTVADYLKFKDLILRMLDYDPKTRIQPYYALQHSFFK    480
NZO/HlLtJ_MGP_NZOHlLtJ_P0045778
RKLHNILGVETGGPGGRRAGESGHTVADYLKFKDLILRMLDYDPKTRIQPYYALQHSFFK    480
NOD/ShiLtJ_MGP_NODShiLtJ_P0044977
RKLHNILGVETGGPGGRRAGESGHTVADYLKFKDLILRMLDYDPKTRIQPYYALQHSFFK    480
PWK/PhJ_MGP_PWKPhJ_P0044675
RKLHNILGVETGGPGGRRAGESGHTVADYLKFKDLILRMLDYDPKTRIQPYYALQHSFFK    480
A/J_MGP_AJ_P0045265
RKLHNILGVETGGPGGRRAGESGHTVADYLKFKDLILRMLDYDPKTRIQPYYALQHSFFK    480
WSB/EiJ_MGP_WSBEiJ_P0044346
RKLHNILGVETGGPGGRRAGESGHTVADYLKFKDLILRMLDYDPKTRIQPYYALQHSFFK    480
C57BL/6J_NP_031916.1
RKLHNILGVETGGPGGRRAGESGHTVADYLKFKDLILRMLDYDPKTRIQPYYALQHSFFK    480

************************************************************

CAST/EiJ_MGP_CASTEiJ_P0045101
KTADEGTNTSNSVSTSPAMEQSQSSGTTSSTSSSSGGSSGTSNSGRARSDPTHQHRHSGG    540
129S1/SvImJ_MGP_129S1SvImJ_P0045278
KTADEGTNTSNSVSTSPAMEQSQSSGTTSSTSSSSGGSSGTSNSGRARSDPTHQHRHSGG    540
NZO/HlLtJ_MGP_NZOHlLtJ_P0045778
KTADEGTNTSNSVSTSPAMEQSQSSGTTSSTSSSSGGSSGTSNSGRARSDPTHQHRHSGG    540
```

```
NOD/ShiLtJ_MGP_NODShiLtJ_P0044977
KTADEGTNTSNSVSTSPAMEQSQSSGTTSSTSSSSGGSSGTSNSGRARSDPTHQHRHSGG    540
PWK/PhJ_MGP_PWKPhJ_P0044675
KTADEGTNTSNSVSTSPAMEQSQSSGTTSSTSSSSGGSSGTSNSGRARSDPTHQHRHSGG    540
A/J_MGP_AJ_P0045265
KTADEGTNTSNSVSTSPAMEQSQSSGTTSSTSSSSGGSSGTSNSGRARSDPTHQHRHSGG    540
WSB/EiJ_MGP_WSBEiJ_P0044346
KTADEGTNTSNSVSTSPAMEQSQSSGTTSSTSSSSGGSSGTSNSGRARSDPTHQHRHSGG    540
C57BL/6J_NP_031916.1
KTADEGTNTSNSVSTSPAMEQSQSSGTTSSTSSSSGGSSGTSNSGRARSDPTHQHRHSGG    540

************************************************************

CAST/EiJ_MGP_CASTEiJ_P0045101
HFAAAVQAMDCETHSPQVRQQFPAPLGWSGTEAPTQVTVETHPVQETTFHVAPQQNALHH    600
129S1/SvImJ_MGP_129S1SvImJ_P0045278
HFAAAVQAMDCETHSPQVRQQFPAPLGWSGTEAPTQVTVETHPVQETTFHVAPQQNALHH    600
NZO/HlLtJ_MGP_NZOHlLtJ_P0045778
HFAAAVQAMDCETHSPQVRQQFPAPLGWSGTEAPTQVTVETHPVQETTFHVAPQQNALHH    600
NOD/ShiLtJ_MGP_NODShiLtJ_P0044977
HFAAAVQAMDCETHSPQVRQQFPAPLGWSGTEAPTQVTVETHPVQETTFHVAPQQNALHH    600
PWK/PhJ_MGP_PWKPhJ_P0044675
HFAAAVQAMDCETHSPQVRQQFPAPLGWSGTEAPTQVTVETHPVQETTFHVAPQQNALHH    600
A/J_MGP_AJ_P0045265
HFAAAVQAMDCETHSPQVRQQFPAPLGWSGTEAPTQVTVETHPVQETTFHVAPQQNALHH    600
WSB/EiJ_MGP_WSBEiJ_P0044346
HFAAAVQAMDCETHSPQVRQQFPAPLGWSGTEAPTQVTVETHPVQETTFHVAPQQNALHH    600
C57BL/6J_NP_031916.1
HFAAAVQAMDCETHSPQVRQQFPAPLGWSGTEAPTQVTVETHPVQETTFHVAPQQNALHH    600

************************************************************

CAST/EiJ_MGP_CASTEiJ_P0045101
HHGNSSHHHHHHHHHHHHHHHGQQALGNRTRPRVYNSPTNSSSTQDSMEVGHSHHSMTSLSS    660
129S1/SvImJ_MGP_129S1SvImJ_P0045278
HHGNSSHHHHHHHHHHHHHHHGQQALGNRTRPRVYNSPTNSSSTQDSMEVGHSHHSMTSLSS    660
NZO/HlLtJ_MGP_NZOHlLtJ_P0045778
HHGNSSHHHHHHHHHHHHHHHGQQALGNRTRPRVYNSPTNSSSTQDSMEVGHSHHSMTSLSS    660
NOD/ShiLtJ_MGP_NODShiLtJ_P0044977
HHGNSSHHHHHHHHHHHHHHHGQQALGNRTRPRVYNSPTNSSSTQDSMEVGHSHHSMTSLSS    660
PWK/PhJ_MGP_PWKPhJ_P0044675
HHGNSSHHHHHHHHHHHHHHHGQQALGNRTRPRVYNSPTNSSSTQDSMEVGHSHHSMTSLSS    660
A/J_MGP_AJ_P0045265
HHGNSSHHHHHHHHHHHHHHHGQQALGNRTRPRVYNSPTNSSSTQDSMEVGHSHHSMTSLSS    660
WSB/EiJ_MGP_WSBEiJ_P0044346
HHGNSSHHHHHHHHHHHHHHHGQQALGNRTRPRVYNSPTNSSSTQDSMEVGHSHHSMTSLSS    660
C57BL/6J_NP_031916.1
HHGNSSHHHHHHHHHHHHHHHGQQALGNRTRPRVYNSPTNSSSTQDSMEVGHSHHSMTSLSS    660

************************************************************

CAST/EiJ_MGP_CASTEiJ_P0045101
STTSSSTSSSSTGNQGNQAYQNRPVAANTLDFGQNGAMDVNLTVYSNPRQETGIAGHPTY    720
129S1/SvImJ_MGP_129S1SvImJ_P0045278
STTSSSTSSSSTGNQGNQAYQNRPVAANTLDFGQNGAMDVNLTVYSNPRQETGIAGHPTY    720
NZO/HlLtJ_MGP_NZOHlLtJ_P0045778
STTSSSTSSSSTGNQGNQAYQNRPVAANTLDFGQNGAMDVNLTVYSNPRQETGIAGHPTY    720
NOD/ShiLtJ_MGP_NODShiLtJ_P0044977
STTSSSTSSSSTGNQGNQAYQNRPVAANTLDFGQNGAMDVNLTVYSNPRQETGIAGHPTY    720
PWK/PhJ_MGP_PWKPhJ_P0044675
STTSSSTSSSSTGNQGNQAYQNRPVAANTLDFGQNGAMDVNLTVYSNPRQETGIAGHPTY    720
A/J_MGP_AJ_P0045265
STTSSSTSSSSTGNQGNQAYQNRPVAANTLDFGQNGAMDVNLTVYSNPRQETGIAGHPTY    720
WSB/EiJ_MGP_WSBEiJ_P0044346
STTSSSTSSSSTGNQGNQAYQNRPVAANTLDFGQNGAMDVNLTVYSNPRQETGIAGHPTY    720
C57BL/6J_NP_031916.1
STTSSSTSSSSTGNQGNQAYQNRPVAANTLDFGQNGAMDVNLTVYSNPRQETGIAGHPTY    720

************************************************************

CAST/EiJ_MGP_CASTEiJ_P0045101          QFSANTGPAHYMTEGHLAMRQGADREESPMTGVCVQQSPVASS    763
129S1/SvImJ_MGP_129S1SvImJ_P0045278    QFSANTGPAHYMTEGHLAMRQGADREESPMTGVCVQQSPVASS    763
NZO/HlLtJ_MGP_NZOHlLtJ_P0045778        QFSANTGPAHYMTEGHLAMRQGADREESPMTGVCVQQSPVASS    763
NOD/ShiLtJ_MGP_NODShiLtJ_P0044977      QFSANTGPAHYMTEGHLAMRQGADREESPMTGVCVQQSPVASS    763
PWK/PhJ_MGP_PWKPhJ_P0044675            QFSANTGPAHYMTEGHLAMRQGADREESPMTGVCVQQSPVASS    763
A/J_MGP_AJ_P0045265                    QFSANTGPAHYMTEGHLAMRQGADREESPMTGVCVQQSPVASS    763
WSB/EiJ_MGP_WSBEiJ_P0044346            QFSANTGPAHYMTEGHLAMRQGADREESPMTGVCVQQSPVASS    763
C57BL/6J_NP_031916.1                   QFSANTGPAHYMTEGHLAMRQGADREESPMTGVCVQQSPVASS    763
                                       *******************************************
```

Table S1. Sequence alignment of Dyrk1a gene from the eight CC founders found no differences in the protein sequence.
